# Decoder-FFPE-seq enables sensitive, genome-wide spatial transcriptomics of archival tissues at single-cell resolution

**DOI:** 10.1101/2025.09.12.675967

**Authors:** Jiao Cao, Jun Zeng, Yiwei Meng, Shanqing Huang, Jing Zhang, Xing Fang, Yang Guo, Weixiong Shi, Yutong Liu, Zhong Zheng, Jianchen Fang, Xuepeng Xiong, Gang Chen, Lingling Wu, Chaoyong Yang

## Abstract

Formalin-fixed paraffin-embedded (FFPE) tissue-compatible spatial transcriptomics (ST) is transforming biology and histopathology, but achieving high sensitivity, genome-wide coverage, single-cell resolution and researcher-accessibility remains challenging. Here, we present a dendrimeric DNA coordinate-barcoding design for FFPE-adapted spatial RNA sequencing (Decoder-FFPE-seq), by integrating RNA-to-DNA probe conversion with blotting-assisted barcoding. Decoder-FFPE-seq provides single-cell resolution with a 2.54-4.25-fold sensitivity increase in mouse brain transcriptomic profiling over similar assays with an available protocol, successfully mapping rare genes and sparse cell types. It robustly resolves tumor microenvironments (TMEs) in clinical archives preserved 6-20 years. With expanded probe libraries, Decoder-FFPE-seq captures long non-coding and viral RNAs, revealing spatial co-expression networks with mRNAs in oral squamous cell carcinomas (OSCCs). In an OSCC immunotherapy cohort, it delineates TME heterogeneity across pathology response groups and tracks spatial cellular dynamics pre- and post-immuno-chemotherapy. Decoder-FFPE-seq establishes a robust, high-resolution framework for sensitive ST profiling in fundamental and translational studies of archival tissues.

## Main

Spatial transcriptomics (ST) uncovers cellular architecture and molecular networks within native tissue context, advancing insights into development biology, organogenesis, and disease pathology^1–5^. In oncology, ST provides unprecedented resolution to deconstruct the tumor microenvironment (TME), revealing its cellular composition, spatial organization, and interactions, unraveling the underlying molecular mechanisms in tumor initiation, progression, and therapy resistance^6, 7^. Translating these insights into clinical practice requires ST platforms compatible with formalin-fixed paraffin-embedded (FFPE) samples, the foundation of pathology archives^8^. These vast repositories offer unparalleled opportunities to link spatial molecular landscapes with annotated clinical outcomes, thereby driving precision oncology. However, RNAs within FFPE samples undergo crosslinking, fragmentation, and degradation during fixation and long-term storage. These alterations induce poly(A) tail loss and adduct formation between nucleic acids and biomolecules^9^, undermining poly(T)-based RNA capture and enzymatic reactions^10^, hampering the ST application to archival specimens.

Effective ST profiling in FFPE samples requires four key capabilities: (1) genome-wide recovery of low-quality transcripts, spanning mRNAs, regulatory RNAs, and pathogenic RNAs, to reconstruct accurate spatial gene-expression and co-regulatory networks; (2) sensitive detection of low-abundance or severely degraded transcripts in archived tissues, enabling biomarker identification and mechanistic insight into disease processes; (3) cellular-resolution mapping of structural details and cell–cell interactions; and (4) accessibility for widespread use across research settings to enable longitudinal cohort studies. Despite several pioneer explorations, options for FFPE-compatible ST methods that satisfy these requirements are limited.

Imaging-based ST methods such as multiplexed fluorescence *in situ* hybridization (FISH) and *in situ* sequencing platforms (e.g., 10× Genomics’ Xenium^11^, NanoString’s CosMx^12, 13^, Vizgen’s MERSCOPE^14^) achieve subcellular resolution and high sensitivity in FFPE tissues^15^, but their limited throughput, labor-intensive workflows, and imaging-related signal drift preclude transcriptome-wide scalability. In contrast, spatially barcoded array-based sequencing methods such as BGI’s

Stereo-seqV2^16^ and Patho-DBiT^17^ achieve high-throughput whole-transcriptome sequencing via *in situ* capture and spatial barcoding using random primers or *in situ* polyadenylation. However, they suffer reduced sensitivity due to ribosomal RNA (∼80%) dominance^18^ and inefficient enzymatic reactions in heavily cross-linked FFPE tissues. Additionally, Patho-DBiT is hindered by tissue deformation and cross-contamination risks across microchannels, while Stereo-seqV2 requires complex position decoding, both of which restrict accessibility.

We previously developed Decoder-seq^19^, employing pioneering dendrimeric DNA coordinate barcoding slides (Decoder slides) containing high-density deterministic spatial barcodes for sensitive ST analysis in fresh-frozen (FF) tissues. Here we report FFPE tissue-adapted Decoder-seq (Decoder-FFPE-seq), which enables sensitive, high-resolution, whole-transcriptome profiling of archival tissues in a widely accessible manner. This is achieved by converting RNAs into poly(A)-tailed DNA probes, followed by blotting-assisted spatial barcoding and sequencing. Unlike commercial Visium CytAssist_FFPE^11, 20^/Visium HD^21^ platforms (10× Genomics), which also adopt probe-targeted strategies but are limited to mRNA profiling, relatively low sensitivity, and closed protocols, Decoder-FFPE-seq incorporates several major advances. First, we broadened the scope of ST to wider RNA species to deepen our understanding of spatial biology. Multiple split-probe libraries were designed for specific and unbiased recovery of whole mRNAs, key regulatory long non-coding RNAs (lncRNAs), and pathogenic RNAs. Second, we implemented a blotting-assisted spatial barcoding strategy, which separates tissue pretreatment and probe labeling on independent slides, thereby preventing barcode loss and exploiting Decoder slides’ high-density barcodes for enhanced sensitivity. Third, Decoder-FFPE-seq achieves single-cell resolution (8 μm spots), allowing for high-definition gene and cell profiling. Fourth, we developed a standardized workflow, spanning tissue pretreatment, probe targeting, and blotting-assisted spatial barcoding to maximize RNA recovery while ensuring reproducibility and accessibility.

With these innovations, Decoder-FFPE-seq provides enhanced gene detection sensitivity than Decoder-seq and similar FFPE-compatible ST platforms when using our optimized protocol. Compared with Visium HD and Stereo-seq V2, it achieves a 2.54–4.25-fold sensitivity increase in mouse brain profiling at single-cell resolution (8 μm spots), effectively detects low-abundance non-sensory G protein–coupled receptors (GPCRs) and resolves sparse inhibitory neuron subtypes in fine-grained tissue structures. Our approach consistently exhibits great potential in unlocking the spatial transcriptomes of challenging long-term preserved archival tissues dating back 6-20 years, even reliably resolves the TME of a 20-year-old oral squamous cell carcinoma (OSCC) with a distribution value at 200 nucleotides (DV200) of only 17%. By simultaneously performing spatial co-profiling of mRNA-lncRNA and host mRNA–pathogenic RNA, Decoder-FFPE-seq uncovers their interaction networks that regulate tumor progression. Application to an immune checkpoint blockade (ICB) trial of OSCC samples, it reveals spatial TME heterogeneity across pathology response groups and tracks spatiotemporal dynamics of key cellular and molecular compartments underlying different responses to immuno-chemotherapy. Collectively, Decoder-FFPE-seq provides a powerful and accessible platform for single-cell, genome-wide spatial transcriptomics in archival tissues, opening new avenues for both fundamental biology and translational research.

## Results

### Workflow of Decoder-FFPE-seq

Decoder-FFPE-seq consists of three main steps: (i) RNA-to-DNA probe conversion, (ii) blotting-assisted spatial barcoding, and (iii) sequencing and bioinformatic analysis (**Fig. 1a**). To recover crosslinked, fragmented, and degraded mRNAs, lncRNAs, and pathogenic RNAs from FFPE tissue, we designed split-probe panels that hybridize to each transcript at one or more adjacent sites. Each left probe (LP) carries a PCR handle (Read 2) and a 25-nt targeting sequence, while the right probe (RP) contains a 5′-phosphorylated 25-nt targeting sequence and a poly(A) tail (Methods, **Fig. 1a, i and Supplementary Fig. 1a**). Only correctly paired probes hybridizing adjacently are ligated into a complete sequence bearing both the 3′ poly(A) tail and 5′ Read 2, enabling subsequent spatial barcoding and PCR amplification. This split-probe design eliminates nonspecific binding of single probes, ensuring highly specific and accurate RNA detection in Decoder-FFPE-seq.

**Fig. 1.**
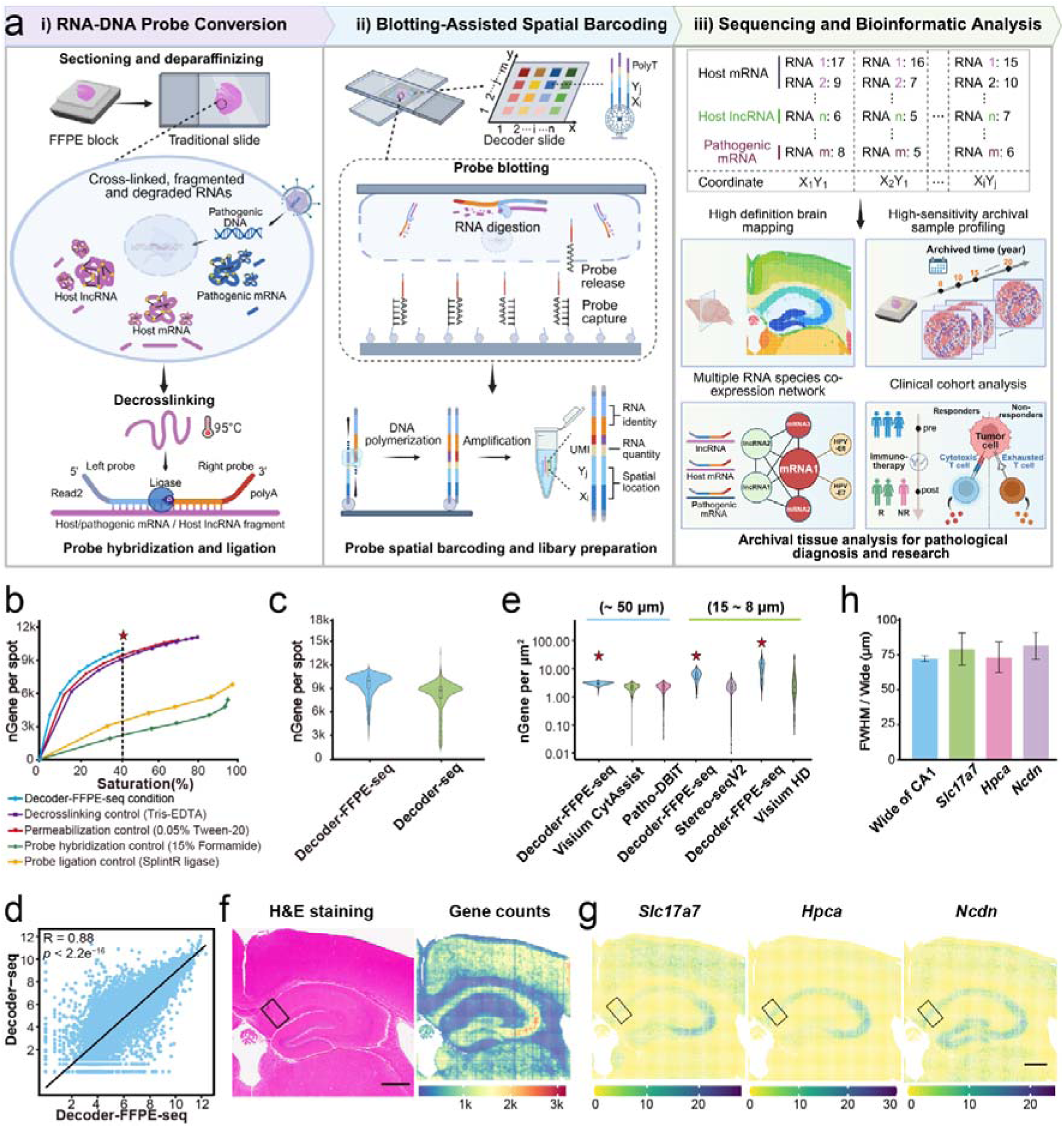
Workflow and performance of Decoder-FFPE-seq in mouse tissues. **(a)** Schematic workflow. (i) RNA-to-DNA probe conversion. FFPE sections are processed by de-paraffinization, decrosslinking, and permeabilization, followed by probe hybridization and ligation. (ii) Blotting-assisted spatial barcoding. Tissue is aligned with barcoded DNA array. Ligated probes are released, captured, and extended with UMIs, spatial barcodes, and PCR handles for NGS. (iii) Sequencing and bioinformatics analysis. Probe sequences, UMIs, and spatial barcodes are extracted to decode RNA identity, abundance, and location for applications of mouse models, clinically archival samples, and ICB-treated clinical cohorts. (**b**) Median genes per spot across conditions tested for decrosslinking, permeabilization, probe hybridization, and ligation in MOBs. (**c**) Comparisons of the number of median genes detected in the Decoder-FFPE-seq and Decoder-seq datasets. (**d**) Gene-gene scatter plots between Decoder-FFPE-seq data (x-axis) and Decoder-seq data (y-axis) of log1p-transformed UMI counts. (**e**) Distribution of gene counts per μm² from FFPE mouse brains of these methods. In Patho-DBiT, only mRNA counts were used. Red stars denote Decoder-FFPE-seq’s performance. (**f**) H&E staining image (left) and gene (right) count maps of the mouse brain using 8 μm spots. (**g**) Expression of CA1 marker genes in the mouse hippocampus. Black boxes indicate regions selected for diffusion analysis. (**h**) Comparison of actual CA1 width versus FWHM of marker gene expression. (**Fig. 1g; Supplementary Fig. 6**).

We developed a blotting-assisted strategy to transfer ligated probes onto Decoder slides for spatial barcoding (**Fig. 1a, ii and Supplementary Fig. 1b**). Using a custom-built alignment instrument, the tissue region of interest is precisely matched with the Decoder slide’s DNA barcode array. Upon RNA digestion and tissue removal, ligated probes are released, captured by the barcoded DNAs, and enzymatically extended to incorporate unique molecular identifiers (UMIs), spatial barcodes, and Read 1 for amplification, library preparation, and sequencing (**Supplementary Fig. 1c-d**). This blotting strategy attaches tissues onto traditional slides rather than Decoder slides, which avoids spatially barcoded DNAs loss from high-temperature treatment of tissue baking and de-crosslinking, and prevents poly(A) tailing-probe blocking (**Supplementary Fig. 2**), enhancing gene detection sensitivity. With next generation sequencing (NGS) and bioinformatics analysis, RNAs were remapped to their original spatial coordinates, achieving mouse brain architecture reconstruction, TME profiling in long-term archival samples, spatial RNA co-regulatory network discovery in OSCCs, and spatial cellular dynamics tracking in clinical cohorts undergoing ICB therapy (**Fig. 1a, iii**).

To optimize transcript recovery from FFPE tissues, we systematically evaluated decrosslinking, permeabilization, probe hybridization, and ligation conditions using mouse olfactory bulbs (MOBs) as a model (**Methods**). The optimized Decoder-FFPE-seq protocol, combined with a 21,178 split-probe panel covering the entire mouse protein-coding transcriptome (**Supplementary Table 1**), detected ∼19,297 unique genes with a median of 60,466 UMIs and 9,960 genes per 50 μm spot at approximately 40% sequencing saturation (**Fig. 1b and Supplementary Fig. 3a**). This demonstrated a substantial improvement in gene detection compared to control treatment conditions of conventional FISH or commercial 10× Genomics with sensitivity even higher than our published Decoder-seq data on FF MOBs (**Supplementary Table 2 and Fig. 1c**). Moreover, the protein-coding gene counts exhibited a high concordance between the Decoder-FFPE-seq and Decoder-seq datasets (R = 0.88, *p* < 2.2e^−16^, **Fig. 1d and Supplementary Fig. 3b**), accompanied by consistent spatial distributions of layer-specific marker genes (**Supplementary Fig. 3c**). Unsupervised clustering faithfully recapitulated the layered MOB structure (**Supplementary Fig. 3d**). These results confirmed that Decoder-FFPE-seq achieves FFPE tissue spatial profiling with performance comparable to FF tissue. To facilitate adoption, we provide a step-by-step protocol in the **Supplementary Information**, designed for implementation in standard laboratories using readily available reagents and equipment.

### Evaluation and benchmarking of Decoder-FFPE-seq

We further benchmarked the performance of Decoder-FFPE-seq against other FFPE-compatible ST platforms, including Visium CytAssist_FFPE, Patho-DBiT, Stereo-seqV2, and Visium HD (**Fig. 1e**). 50 μm-spot Decoder-FFPE-seq detected a median of 29,907 UMIs and 7,571 genes per spot in the mouse brain tissue, outperforming both 55 μm-spot Visium CytAssist_FFPE and 50 μm-pixel Patho-DBiT (**Supplementary Fig. 4a-c**). After normalizing UMI and gene counts across sequencing depths, Decoder-FFPE-seq still show superior sensitivity (**Supplementary Fig. 4d**). Reproducibility between replicates was high (R = 0.99, *P* < 2.2×10^−1^□, **Supplementary Fig. 4e**), and region-specific marker genes were robustly detected and spatially mapped, faithfully reflecting the corresponding anatomical domains (**Supplementary Fig. 4f-g**).

To enable finer gene mapping and high-definition cellular profiling, we improved the spatial resolution of Decoder-FFPE-seq to near-cellular level (15 μm and 8 μm spots). Despite reduced spot size, Decoder-FFPE-seq retained high sensitivity, detecting 1,328 genes per spot at 15 μm and 617 genes per spot at 8 μm (**Fig. 1e-f and Supplementary Fig. 5a**). These results substantially exceeded comparable-resolution methods, such as Stereo-seqV2 (10 μm bins: 232 genes) and Visium HD (8 μm bins: 145 genes). When normalized per unit area (μm²), Decoder-FFPE-seq achieved 2.54–4.25-fold higher gene detection sensitivity than the two methods (**Fig. 1e**). Notably, under low sequencing depth (80 M reads), 8 μm-spot Decoder-FFPE-seq delivered a striking 7.46–27.71-fold improvement in sensitivity (nGene per μm²: Decoder-FFPE-seq, 9.70; Visium HD, 1.30; Stereo-seqV2, 0.35), underscoring its potential to detect low-abundance transcripts (**Supplementary Fig. 5b-c**). Replication of the datasets again showed high concordance (R = 0.98, *P* < 2.2 × 10^−1^□) (**Supplementary Fig. 5d**).

Spatial accuracy of RNA detection, another key quality parameter for high-resolution ST, was evaluated at 8 μm resolution. Focusing on the hippocampal cornu ammonis 1 (CA1) region which contains a densely packed pyramidal cell layer, we assessed lateral mRNA diffusion by calculating the full-width at half-maximum (FWHM) of the expression plots of three region-enriched genes (*Slc17a7*, *Hpca*, and *Ncdn*) (**Fig. 1g and Methods**). Their FWHM values closely matched the actual width of corresponding CA1 regions observed in H&E staining, confirming the well-controlled diffusion of Decoder-FFPE-seq (**Fig. 1h and Supplementary Fig. 6a**). Compared with similar ST platforms, Decoder-FFPE-seq showed a comparable capacity for RNA diffusion control (**Supplementary Fig. 6b-e**). Collectively, these data demonstrate the high sensitivity and spatial accuracy of Decoder-FFPE-seq for ST profiling.

### Decoder-FFPE-seq profiles the mouse brain with high sensitivity and accuracy at single-cell resolution

We next applied 8 μm-spot Decoder-FFPE-seq to profile low-abundance genes, fine structural details, and sparse neuronal subtypes in the mouse brain. Non-sensory GPCRs (∼300-400 genes) are all expressed in brain tissues, play essential roles in brain development, neural signaling, immunity, and metabolism, but are difficult to profile due to low expression levels^22^. Decoder-FFPE-seq successfully detected 340 non-sensory GPCRs (**Supplementary Table 3**), with UMI counts per μm² markedly higher than Visium HD and Stereo-seqV2 (**Fig. 2a and Supplementary Tables 4 –5**).

**Fig. 2.**
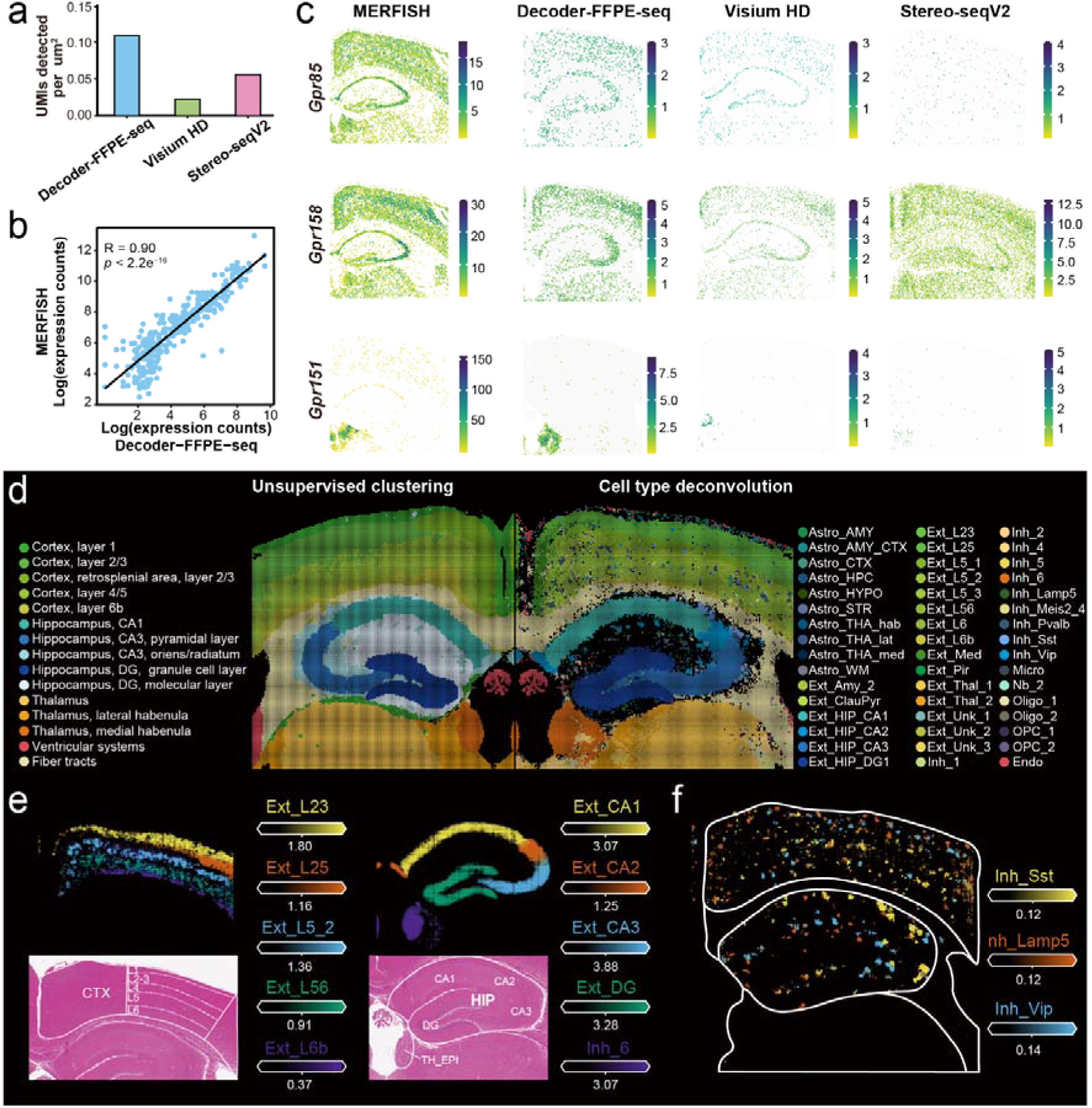
Decoder-FFPE-seq profiles the mouse brain with high sensitivity and accuracy at single-cell resolution. **(a)** Non-sensory GPCR UMIs detected per μm² via 8 μm-spot Decoder-FFPE-seq, Visium HD, and Stereo-seqV2. (**b**) Correlation of GPCR expression between 8 μm-spot Decoder-FFPE-seq and MERFISH data. (**c**) Spatial distribution of representative GPCR genes (*Gpr151*, *Gpr85*, *Gpr158*) across platforms. (**d**) Unsupervised clustering of spots (left) and spatial visualization of deconvolved cell types (right) from the mouse hemibrain section by Decoder-FFPE-seq. (**e**) Fine-grained cortical layers and hippocampal subdivisions. (**f**) Detection and localization of sparse inhibitory neurons.

Its non-sensory GPCR expression displayed a stronger correlation with high-sensitivity MERFISH (R = 0.90, *P* < 2.2×10^−1^□) than the two other methods (**Fig. 2b and Supplementary Fig. 7a**). Spatial distributions of representative genes *Gpr151*, *Gpr85*, and *Gpr158* closely mirrored MERFISH patterns and were more clearly resolved than in Stereo-seqV2 and Visium HD datasets (**Fig. 2c**). Gene Ontology (GO) analysis showed expected enrichment of biological processes including development, neural excitation, immunity, metabolism, and behavior (**Supplementary Fig. 7b**).

Using a single-nucleus RNA sequencing (snRNA-seq) reference for cell-type deconvolution, 82.5% of spots were assigned to a single cell type, confirming the cellular resolution of Decoder-FFPE-seq (**Supplementary Fig. 7c**). With improved resolution, we observed a finer structural detail of the brain which recapitulated its anatomical morphology (**Fig. 2d**). For example, spatial mapping reconstructed canonical brain architectures, including six cortical layers and hippocampal subregions (CA1, CA2, CA3, dentate gyrus (DG) and the habenula) (**Fig. 2e**).

Subregional subtype distributions matched the spatial expression of marker genes (**Supplementary Fig. 7d-e**). Importantly, rare inhibitory neuron subtypes were also effectively identified in these fine-grained tissue structures (**Fig. 2f**). We observed that Lamp5 and Vip interneurons were enriched in the superficial cortex, while Sst neurons localized to deeper cortical layers and the CA1 stratum oriens. Their distributions were consistent with classical marker expression and prior studies^23^ (**Supplementary Fig. 7f**). Replicate datasets confirmed reproducibility in deconvolution results (R = 0.84, *P* < 2.2×10^−1^□, **Supplementary Fig. 7g**). In summary, 8 μm-spot Decoder-FFPE-seq enables sensitive detection of lowly expressed genes, resolves fine structural details, and captures sparse cell populations, providing a high-definition spatial atlas of FFPE mouse brain architecture.

### Decoder-FFPE-seq characterizes complex TME in long-term archived clinical tissues

To evaluate clinical applicability, Decoder-FFPE-seq was applied to diverse archived tissues using human whole-transcriptome probe panels (**Fig. 3a, Supplementary Fig. 8a–c, and Supplementary Tables 6**). In FFPE breast cancer sections, 50 μm-spot Decoder-FFPE-seq outperformed 55 μm-spot Visium CytAssist_FFPE in UMI and gene counts detected per spot. Its sensitivity in breast cancer and lymph node FFPE tissues even exceeded that of Visium for FF samples. Increasing probe multiplicity from average one to three pairs per transcript enhanced lymph node detection from 30,140 UMIs/7,770 genes per spot to 103,501 UMIs/12,971 genes (**Supplementary Fig. 8b**), demonstrating that sensitivity can be tuned via multi-region probe targeting. Notably, Decoder-FFPE-seq detected a median of 10,625 UMIs and 3,951 genes per spot in a six-year-old OSCC FFPE sample using a single probe-pair design (**Fig. 3a, Supplementary Fig. 8c**).

**Fig. 3.**
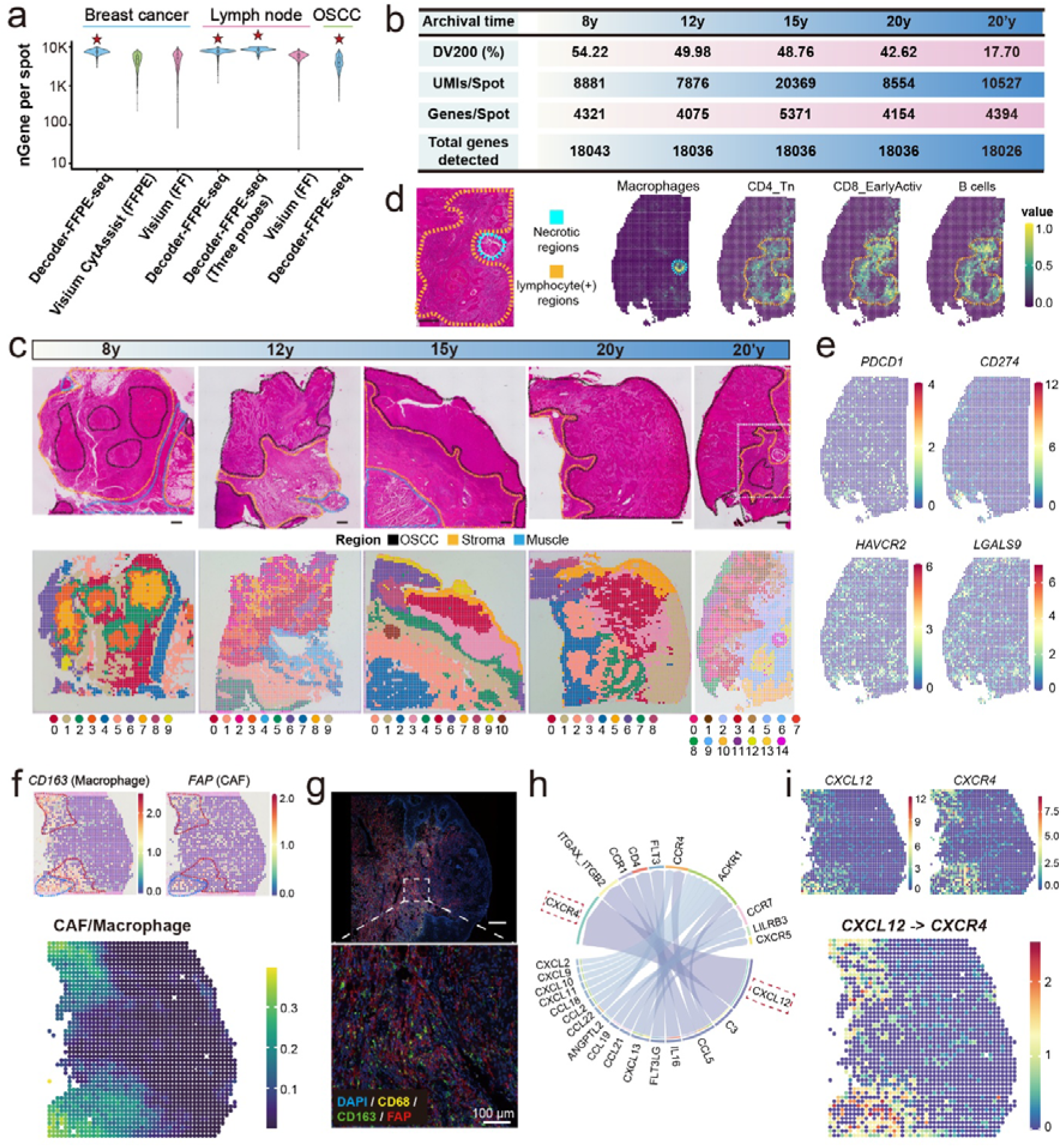
Decoder-FFPE-seq characterizes complex TME in long-term archived clinical tissues. **(a)** Gene counts per spot in FFPE breast cancer, lymph node, and OSCC tissues profiled by Decoder-FFPE-seq, compared with FFPE and FF breast cancer samples profiled by Visium CytAssist_FFPE and Visium. Red stars indicate Decoder-FFPE-seq performance. (**b**) Summary table of archival time, DV200, median gene/UMI counts per spot, and total detected genes for long-term OSCC archives. (**c**) H&E images with pathological annotation (upper) of tumor (black), stroma (yellow) and muscle (blue), and unsupervised clustering of Decoder-FFPE-seq spots from OSCC archives. (**d**) Zoomed H&E image of 20-year-old OSCC with annotated a necrotic area (blue) and lymphocyte aggregates (yellow) and their corresponding cell distribution. **(e)** Spatial distributions of receptors and ligands (**f**) Co-localization of CAFs and macrophages with marker genes. (**g**) Multiplex immunofluorescence of macrophages (CD68/CD163) and CAFs (FAP); DAPI (blue), CD68 (yellow), CD163 (green), FAP (red), right, magnified view. (**h**) Ligand–receptor interactions in cluster 10. **(i)** Spatial expression of CXCL12 and CXCR4 with their interaction intensity in the OSCC section.

Further extending the analysis to 8–20-year archived OSCC specimens, our approach consistently detected a median of over 4,000 genes per spot, even in a 20-year sample with DV200 of 17% (**Fig. 3b and Supplementary Fig. 8d**). Spatial clusters from the unsupervised clustering of these samples closely matched H&E-annotated pathological regions, and cell-type deconvolution confirmed cluster identities (**Fig. 3c and Supplementary Fig. 8e**). For example, in the 20-year OSCC sample with DV200 of 17%, cluster 14 was localized to a necrotic tumor area, showing macrophage distribution, consistent with studies that the tumor-associated macrophage is enriched in hypoxic and necrotic TME^24^ (**Fig. 3d**). While clusters 2, 5, 6, and 8 were confined to lymphocyte (+) stromal regions, where diverse immune populations, including CD4⁺ T cells, CD8⁺ T cells, B cells, and plasma cells, were accurately resolved, corresponding to lymphocyte aggregates on the H&E-stained section (**Fig. 3d**). Besides, we evaluated six housekeeping genes (*UBC*, *PPIB*, *HRPT1*, *TPB*, *GUSB*, *ALAS1*) in this sample. Their expression level from pseudo-bulk transcript counts of Decoder-FFPE-seq data aligned with bulk RNA-seq from TCGA-Head and Neck Squamous Cell Carcinoma (HNSC) cohort, and exhibited expected uniformity of spatial distribution (**Supplementary Fig. 8f–g and Methods**), indicating that RNA degradation during extremely long-term preservation did not distort spatial expression patterns of Decoder-FFPE-seq. Low-abundance immune checkpoint genes were also effectively detected, including classic receptors (*PDCD1*, *HAVCR2*) and ligands (*CD274*, *LGALS9*) (**Fig. 3e**). These results demonstrate Decoder-FFPE-seq’s robustness in unlocking spatial transcriptomes of challenging archival tissues, enabling longitudinal patient cohort studies.

Focusing on the six-year-old OSCC specimen with lymph node metastasis at initial diagnosis, we meticulously explored its intricate TME. Likewise, unsupervised clustering identified 13 clusters, whose spatial projections aligned with pathologically annotated tumor, stroma, muscle, and normal mucosa regions (**Supplementary Fig. 9a**). With cell-type deconvolution and marker gene expression, we intriguingly observed that *CD163*⁺ macrophages co-localized with *FAP*⁺ CAFs, which of both were known to promote tumor progression^25, 26^ (**Fig. 3f**). Multiplex immunofluorescence confirmed their spatial distribution (**Fig. 3g**). Co-localization analysis revealed macrophages predominantly associated with CAFs in cluster 5 (muscle) and cluster 10 (stroma) (**Supplementary Fig. 9b**). Notably, cluster 10, located at the tumor-stroma interface, was enriched with diverse immune cells with upregulation of immune-related genes (**Supplementary Fig. 9c-d**). Enrichment of hallmark gene set highlighted the activation of epithelial-mesenchymal transition (EMT), interferon-gamma response, and allograft rejection pathways (**Supplementary Fig. 9e**), suggesting an immune–tumor antagonistic invasive front, where CAF–macrophage interactions potentially transforming the TME with immunosuppression, tumor promotion, and drug resistance characteristics. Quantification of cell-type percentages confirmed CAFs were the most abundant, accompanied by macrophages (especially M2-like, CD163^+^), monocytes, and regulatory T cells (Tregs) as predominant immune cells (**Supplementary Fig. 9f**).

Ligand–receptor analysis further highlighted dominant CXCL12–CXCR4 signaling pair, whose expression patterns corresponded to macrophage–CAF co-localization (**Fig. 3h-i**). We speculated that the CXCL12-CXCR4 axis may serves as a key driver in shaping a tumor-promoting microenvironment, in which CAF-derived CXCL12 recruits CXCR4⁺ immune cells and induces monocytes differentiation into macrophages, as previously reported^27^. Overall, these findings link spatial cellular architecture to tumor invasion and clinical progression, illustrating Decoder-FFPE-seq’s power for dissecting TME in archived clinical specimens.

### Extending ST to non-poly(A)-tailed and non-host RNAs detection for spatial biology study in archival OSCC samples

Current understanding of transcriptional regulation remains biased toward protein-coding genes, despite the critical contributions of non-coding RNAs and pathogenic RNAs to gene regulation and disease progression^28–30^. Spatial profiling of these RNAs has been hindered by their low abundance, partial lack of polyadenylation, or, in the case of infectious agents, the frequent restriction of samples to FFPE-archived tissues. Leveraging the flexible probe design of Decoder-FFPE-seq, we extended its scope to capture both non-poly(A)-tailed and non-host RNAs, for enabling joint profiling of mRNA–lncRNA and host mRNA–pathogenic RNA of archival OSCC samples (**Fig. 4a**), providing new insights into transcriptional interactions that underlay tumor progression.

**Fig. 4.**
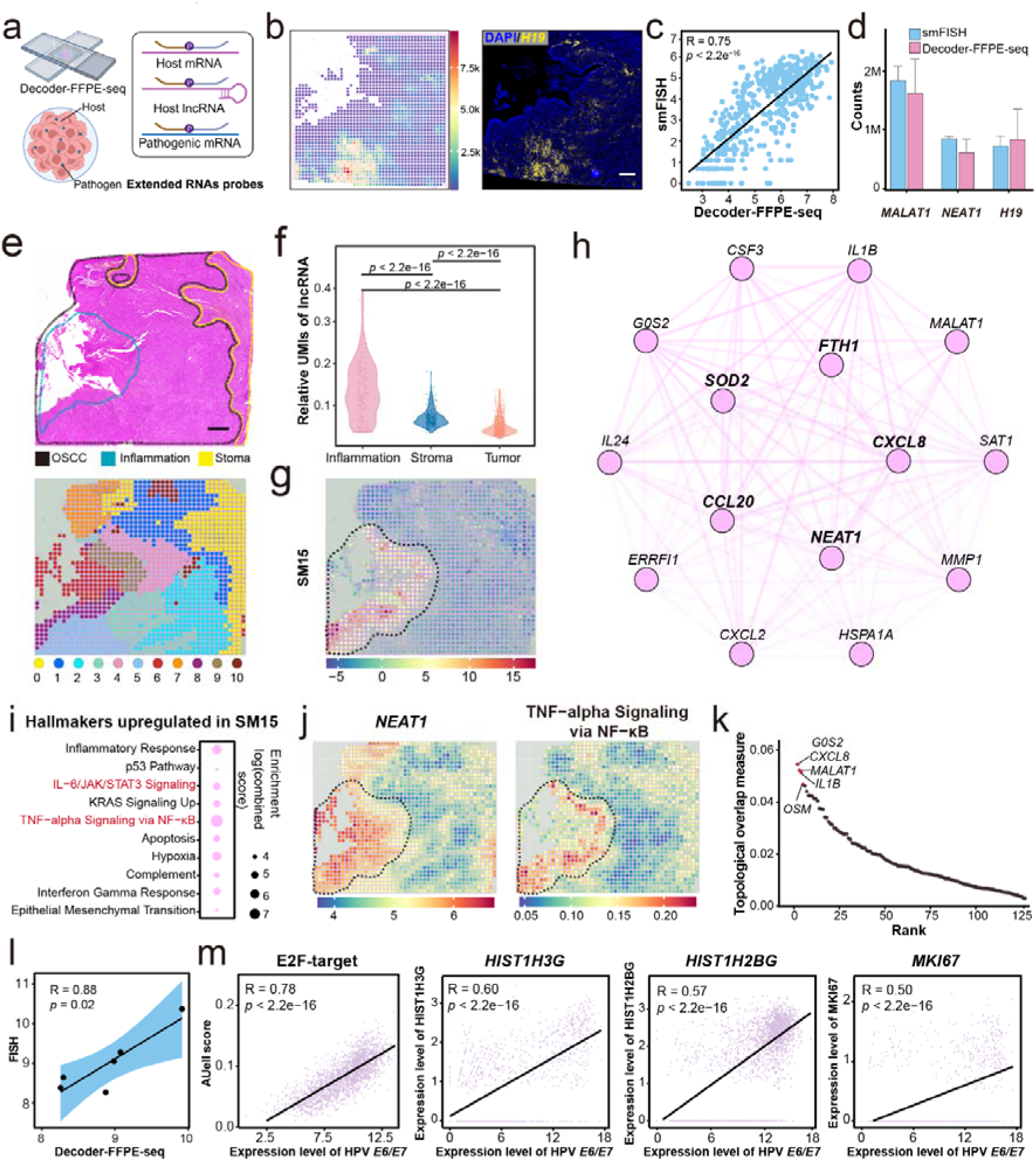
Spatial co-profiling of mRNA-IncRNA and host mRNA-HPV RNA in OSCC archives. **(a)** Schematic of Decoder-FFPE-seq co-profiling for mRNA-IncRNA and host mRNA-HPV RNA. (**b**) Spatial distribution of *H19* from Decoder-FFPE-seq and smFISH in adjacent sections. (**c**) Gene consistency between Decoder-FFPE-seq and smFISH (R = 0.75). (**d**) Comparison of representative lncRNA counts detected by Decoder-FFPE-seq and smFISH (*n* = 3). (**e**) H&E image (top) of OSCC section with pathological annotations (black: OSCC, yellow: stroma, blue: tumor associated-inflammatory region) and its unsupervised clustering (bottom). (**f**) Relative lncRNA expression across annotated regions. (**g**) Spatial distribution of SM15. (**h**) Co-expression network of top 15 hub genes in SM15. (**i**) Hallmark pathways enriched in SM15. (**j**) Spatial distributions of TNFα–NFκB signaling (left) and *NEAT1* expression (right). (**k**) TOM similarity with *NEAT1* in SM15. (**l**) Correlation of HPV transcript counts between Decoder-FFPE-seq and HPV-FISH (*n* = 6 matched regions). (**m**) Spatial correlation of E2F pathway activity and downstream gene expression with HPV expression.

LncRNAs exhibit tissue- and cancer-specific expression and regulate mRNA to promote oncogenesis through diverse mechanisms^31, 32^. We further designed a probe panel targeting 275 OSCC-associated lncRNAs (**Methods and Supplementary Table 8**). Using the lncRNA-only panel, Decoder-FFPE-seq detected all 275 targets, with a median of 3,358 UMIs and 114 genes per 50-μm spot (**Supplementary Fig. 10a**). LncRNA expression counts reasonably correlated with bulk RNA-seq data from the TCGA-HNSC cohort (R = 0.55, p < 2.2×10^-16^) (**Supplementary Fig. 10b**).

Representative lncRNAs (*H19*, *NEAT1*, *MALAT1*) showed consistent spatial distribution with those measured by gold-standard single molecule FISH (smFISH) in adjacent OSCC sections, exhibiting correlation coefficients exceeding 0.50 (**Fig. 4b–c and Supplementary Fig. 10c**). Quantitative comparisons of three gene counts showed that Decoder-FFPE-seq recovered 92.0 ± 17.1% of numbers detected by smFISH (**Fig. 4d and Supplementary Table 10**), confirming its high sensitivity.

We next explored the regulatory roles of mRNA–lncRNA interactions in OSCC using a combined mRNA–lncRNA panel (**Supplementary Fig. 10d**). Unsupervised clustering identified 11 transcriptionally distinct clusters corresponding to pathologically annotated tumor, stroma, and tumor-associated inflammatory regions (**Fig. 4e**), where these regions showed distinct marker gene expression (**Supplementary Fig. 10e**). Differential expression analysis revealed upregulated and downregulated lncRNAs in tumor versus non-tumor regions (**Supplementary Fig. 10f**). For example, oncogenic lncRNA *H19* and tumor suppressor MEG3 were upregulated and downregulated in tumors, respectively, consistent with their established role^33, 34^ (**Supplementary Fig. 10f-g**). Notably, relative lncRNA expression was significantly enriched in the tumor-associated inflammatory region over stromal or other tumor compartments (**Fig. 4f**), potentially indicating regulatory roles of lncRNAs in such TME. Co-expression network analysis with hdWGCNA identified 40 spatial modules (SMs), of which SM1 and SM15 contained multiple lncRNAs as hub genes with high connectivity with other transcripts (**Supplementary Fig. 11a–c**). Specifically, SM15 was localized to the tumor-associated inflammatory region, and contained hub genes such as the oncogenic lncRNA *NEAT1*, inflammatory mediators *CXCL8* and *CCL20*, and oxidative stress regulators *FTH1* and *SOD2* (**Fig. 4g–h**). Functional enrichment confirmed that SM15 was strongly associated with inflammatory pathways, including TNFα–NFκB, Inflammatory response, and IL-6/JAK/STAT3 (**Fig. 4i**). Spatial projection of TNFα–NFκB pathway was overlapped with *NEAT1* expression, yielding correlation coefficients of 0.63 (**Fig. 4j**), suggesting that *NEAT1* may involve in this signaling axis. Besides, spatial co-expression network analysis revealed highest topological similarity of *NEAT1* with inflammation-related genes (*IL1B*, *CXCL8*) and the cell cycle regulator *G0S2* (**Fig. 4k**). Their co-localization within SM15 supports potential regulatory interactions consistent with prior studies (**Supplementary Fig. 11d**). These findings suggest that *NEAT1* may serve as a hub gene in the co-expression network of mRNA-IncRNA, amplify TNFα signaling to drive inflammatory gene expression (e.g., *IL1B*, *CXCL8*), and coordinate inflammatory responses and tumor progression^35, 36^.

High-risk HPV16/18 are well-established drivers of OSCC through expression of E6/E7 oncoproteins^37^. We designed HPV16/18-specific E6/E7 probes for co-profiling host transcriptome and viral E6/E7 transcripts in HPV-positive OSCC to investigate how pathogen RNAs influence host genes (**Fig. 4a, Methods, and Supplementary Table 8**). Spatial expression of E6/E7 transcripts was confined to tumor regions (**Supplementary Fig. 11e**), and their UMI counts strongly correlated with FISH measurements (R = 0.88, *p* = 0.02; **Fig. 4l**), validating accuracy of Decoder-FFPE-seq in profiling pathogen RNA. Unsupervised clustering identified distinct clusters whose spatial projections aligned with tumor or stromal compartments (**Supplementary Fig. 11f**). Functional enrichment revealed that tumor regions exhibited significant upregulation of *HALLMARK_E2F_TARGETS* and *MYC_TARGETS_V1* compared to the stroma (**Supplementary Fig. 11g**). Of note, *E2F* target and its downstream histone- and cell cycle-related genes (*HIST1H3G*, *HIST1H2BG*, *MKI67*), positively correlated with E6/E7 transcript abundance (**Fig. 4m**). These results are consistent with the known roles of E6/E7 proteins in p53 and pRb inactivation, aberrant E2F activation, and uncontrolled cell cycle progression^38^.

Together, these findings demonstrate the capacity of Decoder-FFPE-seq in dissection of multi-layer transcriptional interactions and host–pathogen regulatory networks in clinical archived samples, advancing spatial RNA biology into previously challenging domains.

### Decoder-FFPE-seq identifies pathological response-related spatial niches after immunotherapy in OSCC

ICB has achieved major clinical success in oncology, yet therapeutic responses vary widely, and the determinants of these divergent outcomes remain incompletely understood^39–41^. To explore this, 50-μm-spot Decoder-FFPE-seq were applied to seven post-treatment FFPE specimens from patients who were diagnosed with locally advanced OSCC and enrolled in neoadjuvant anti-PD-1 clinical trials (**Supplementary Fig. 12 and Supplementary Table 11-12**). Based on residual tumor regression, samples were classified as major pathological response (MPR, *n* = 3), pathological partial response (pPR, *n* = 2), and pathological non-response (pNR, *n* = 2) (**Fig. 5a and Supplementary Fig. 12**).

**Fig. 5.**
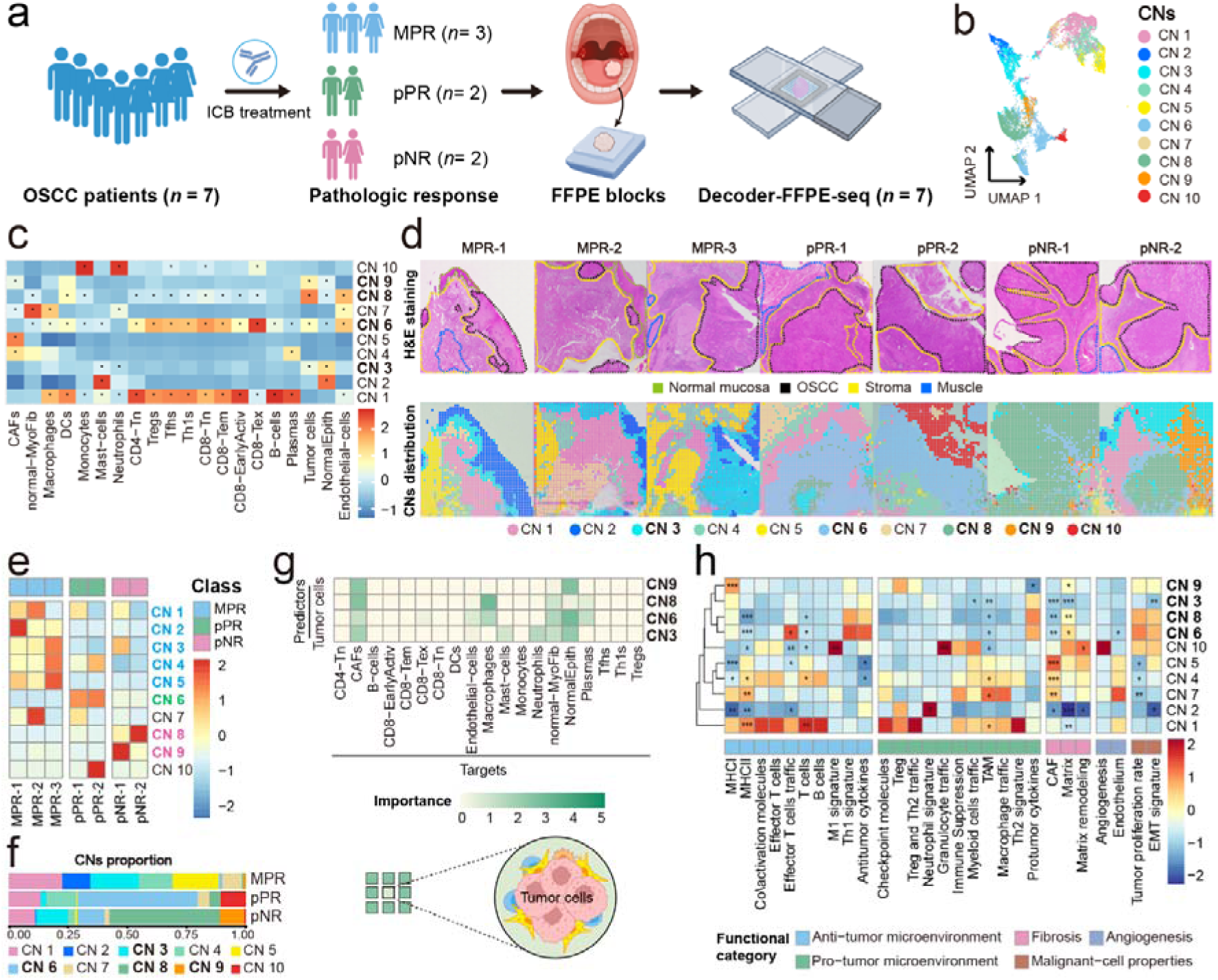
Decoder-FFPE-seq identifies response-related spatial niches after immunotherapy in OSCC. **(a)** Workflow of OSCC sample collection and processing for Decoder-FFPE-seq. (**b**) UMAP of CNs based on deconvolved cell-type compositions. (**c**) Scaled median cell-type composition for each CN. Asterisks indicate significantly increased abundance of a specific cell type compared with other CNs (two-sided Wilcoxon rank-sum test, adjusted *P*L<L0.05). (**d**) H&E images of seven tissue sections (top) with spatial projections of CNs (bottom). Pathologically annotated regions included OSCC (black), stroma (yellow), muscle (light blue), and normal mucosa (green). (**e-f**) Relative CN composition across each patient (**e**) and response groups (**f**). (**g**) Median importance of tumor cell abundance for predicting other cell types within ∼50 μm spots. (**h**) AUCell-based enrichment of tumor-associated pathways. Asterisks indicate pathways significantly upregulated in a given CN compared with others (two-sided Wilcoxon rank-sum test, adjusted *P* < 0.05).

To dissect spatial heterogeneity of the TME across pathology response groups, we applied cell-type deconvolution and unsupervised clustering to Decoder-FFPE-seq data from seven samples, resolving 10 major cell-type niches (CNs) (**Fig. 5b, Supplementary Fig. 13a**). Each CN displayed distinct cell-type compositions and spatial distributions that broadly aligned with pathology-defined regions, suggesting they may represent functional building blocks of the TME (**Fig. 5c–d**). CNs 3, 6, and 8 localized primarily to tumor regions, with CNs 6 and 8 representing immune-infiltrated tumor niches and CN 3 a mixed tumor–epithelium niche enriched for mast cells and neutrophils. Stromal regions contained CNs 1, 4, 5, 7, and 10: CN 1 was an immune-infiltrated stroma, CNs 4, 5, and 7 were CAF-rich, and CN 10 was an inflammation-associated stroma with abundant monocytes and neutrophils. CN 9 represented a CAF-rich tumor–stroma interface niche, while CN 2 corresponded to epithelial regions in normal mucosa (**Fig. 5d**). Notably, niche composition varied across response groups, particularly with tumor cell–enriched CNs (3, 6, 8, and 9) displaying distinct distribution patterns (**Fig. 5e**). MPR tumors were enriched in the mixed tumor–epithelium CN 3, pPR tumors were dominated by CN 6, while pNR tumors contained predominantly tumor-enriched CNs 8 and 9 (**Fig. 5f**).

We next examined functional features of these niches and their association with treatment efficacy. We focused on tumor cell-enriched niches to dissect their heterogeneous spatial interaction patterns between malignant cells and non-neoplastic populations. Using relative tumor cells proportions, the abundances of other cell types were predicted across intra-spot (∼50 μm) (**Fig. 5g**). In MPR groups, the dominant mixed tumor–epithelium CN 3 neighbored epithelial CN 2 and showed strong spatial co-occurrence of tumor cells with CAFs, endothelial cells, mast cells, neutrophils, epithelial cells, and myofibroblasts, suggesting a regression zone with localized inflammation responsive to ICB. In pPR tumors, the immune-infiltrated CN 6 predominated, where tumor cells most strongly predicted CAFs, exhausted CD8^+^ T cells (Tex), endothelial cells, macrophages, myofibroblasts, epithelial cells, and plasma cells. This CD8^+^ Tex-enriched microenvironment likely underlies partial ICB responsiveness. In contrast, pNR tumors were dominated by: (1) CN 8, where tumor cells predicted CAFs, macrophages, myofibroblasts, and plasma cells, reflecting an immunosuppressive TME associated with resistance; and (2) CN 9, where tumor cells co-occurred with CAFs and epithelial cells at the tumor–stroma interface, suggesting roles in ECM remodeling and immune evasion.

Gene set enrichment confirmed niche-specific functional programs (**Fig. 5h**). In MPR tumors, CN 3 showed reduced malignant potential, with low proliferative activity and significantly diminished EMT signatures. In pPR tumors, CN 6 exhibited elevated effector T-cell trafficking, Th1 signaling, and anti-tumor cytokine expression, consistent with immune engagement. Meanwhile, CN 6 upregulated proliferative pathways (E2F, MYC, G2/M cell-cycle progression), TGF-β signaling, and inflammatory programs (**Supplementary Fig. 13b**), reflecting complex tumor–immune crosstalk that may constrain aggressive growth. By contrast, pNR-dominant CN 8 showed elevated pro-tumor cytokines and reduced anti-tumor cytokines, consistent with ineffective immune activity (**Fig. 5h**). GO enrichment of differentially expressed genes between CN 6 and CN 8 further supported this divergence (**Supplementary Fig. 13c**). CN 9 was marked by a strong Treg signature, supporting its role in immunosuppression (**Fig. 5h)**. Stromal niches also exhibited distinct behaviors. CN 1, an immune-infiltrated stroma, upregulated interferon-α/γ responses and displayed partial anti-tumor activity (**Supplementary Fig. 13d**). While both CNs 4 and 5 were CAF-rich, CN 4 facilitated immune infiltration, whereas CN 5 was ECM-enriched and promoted matrix remodeling (**Supplementary Fig. 13e**). The predominance of CN 5 in MPR tumors suggests a “tissue reconstruction” zone undergoing regression, whereas CNs 1 and 4, observed in both MPR and pPR, likely represent immune-activated stromal regions. Collectively, Decoder-FFPE-seq resolves spatial organization of heterogeneous TME with distinct functional states across pathology response groups.

### Decoder-FFPE-seq decodes spatial cellular dynamics in OSCC samples before and after immuno-chemotherapy

Cellular interactions within the TME prior to treatment may govern immune responses to ICB^41, 42^. To elucidate the spatial cellular dynamics of the TME underwent treatment and the corresponding underlying molecular mechanism, we further sequenced the pre-treatment biopsy samples from OSCC patients and performed integrated spatial profiling with matched post-treatment sample data (**Fig. 6a, Supplementary Table 13, and Supplementary Fig. 14**). As different ICB treatments may differentially affect the immune response, paired analyses were limited to samples from patients receiving the same treatment mode (chemo-immunotherapy, MPR groups and pNR-1, **Fig. 6a**). Specifically, the dynamic changes of the TME in paired pre-post treatment FFPE samples (four groups: pre-MPR, post-MPR, pre-pNR-1, and post-pNR-1; n = 8) from 50-μm-spot Decoder-FFPE-seq were investigated, followed by high-definition profiling at single-cell resolution with 8-μm-spot Decoder-FFPE-seq (n = 4) and commercial Xenium (n = 2) on representative samples from four groups (**Fig. 6a**).

**Fig. 6.**
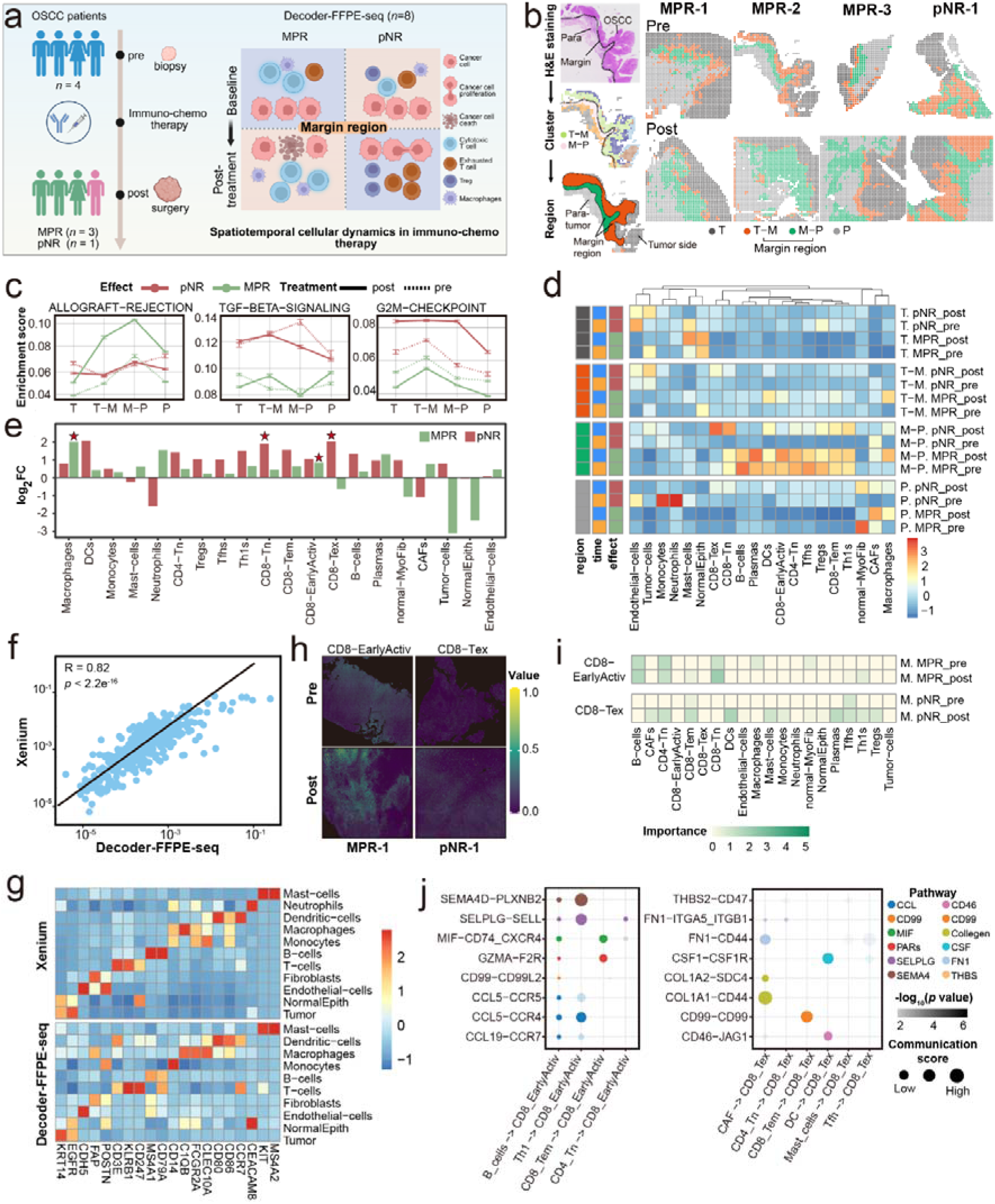
Decoder-FFPE-seq decodes spatial cellular dynamics in OSCC samples before and after immuno-chemotherapy. **(a)** Schematic diagram of sample collection, processing and spatial profiling. **(b)** Schematic diagram of the regional division (left) and spatial distribution of distinct regions in OSCC samples before and after immuno-chemotherapy (right). **(c)** Hallmark pathway dynamic in different spatial regions before and after treatment among the two groups. **(d)** Heatmap of baseline and post-treatment cellular abundance across tissue regions. **(e)** Treatment-induced cellular abundance changes in the M region. **(f)** Comparison of gene expression levels between Decoder-FFPE-seq (UMIs per spot) and Xenium (transcripts per cell). **(g)** Marker gene expression across cell types between Xenium and Decoder-FFPE-seq. **(h)** Spatial distribution of CD8⁺ EarlyActive in MPR-1 and CD8⁺ Tex cells in pNR-1 samples before and after treatment from 8μm-spot Decoder-FFPE-seq. **(i)** Median importance of CD8⁺ EarlyActive in MPR-1 and CD8⁺ Tex in pNR-1 abundance for predicting other cell types within 64-μm radius. **(j)** Cell–cell communication analysis of CD8⁺ EarlyActive (MPR-1) and CD8⁺ Tex (pNR-1) cells with their neighboring cells within the M region.

To systematically characterize dynamic changes in the TME before and after immuno-chemotherapy, we compared the four groups at both molecular and cellular compositional levels across distinct spatial regions, including the tumor (T), tumor-adjacent margin (T-M), margin-adjacent paratumor (M-P), and paratumor (P) zones. These spatial zones were independently delineated in each group via unsupervised clustering and tumor border identified H&E annotation (**Fig. 6b, Supplementary Fig. 15a**). Pathway analysis revealed marked spatial transcriptional heterogeneity among these regions between patient groups (**Fig. 6c, Supplementary Fig. 15b**). By comparing pre- and post–immuno-chemotherapy samples, pathways associated with allograft rejection and IL2/STAT5 signaling were upregulated in MPR groups following treatment, particularly enriched in the T-M or M-P regions. Conversely, these pathways were downregulated in the pNR-1 group, which instead exhibited higher enrichment of TGF-β signaling and proliferative related pathways (E2F targets, G2M checkpoint, MYC targets, and DNA repair). This suggests that the M region of MPR groups was immune activated after treatment, whereas the TME of pNR-1 retained immunosuppressive and proliferative characteristics.

We then calculated the baseline (pre-treatment) cellular abundance and cellular dynamics (defined as the difference between the post-treatment and baseline abundance) based on cell-type deconvolution (**Methods, Supplementary Fig. 15c**). At baseline, the MPR group exhibited pronounced immune cell infiltration in the M-P region, whereas the pNR-1 sample showed a myeloid-dominated microenvironment enriched with monocytes and neutrophils in the P region (**Fig. 6d**). The baseline cellular difference may underlie their divergent therapeutic outcomes, with responders displaying a stronger T cell–mediated immune response that established a favorable anti-tumor front adjacent to the tumor, consistent with observations in pre-immunotherapy colorectal cancer responders^43^ (**Supplementary Fig. 15d**).

Following treatment, the MPR patient exhibited further immune cell enrichment in this region, with the T–M region also showing relatively increased proportions of macrophages, DCs, CD8⁺ EarlyActive. In contrast, the pNR-1 sample showed an elevated proportion of CD8⁺ Tex, CD8⁺/CD4⁺ Tn, DCs, CD8⁺ Tem, Th1, Treg, and macrophages in the M–P region (**Fig. 6d, Supplementary Fig. 15e**). Notably, macrophages showed the greatest increase in the M region of post-MPR samples, with CD8⁺ EarlyActive being the most elevated T cell subset, whereas CD8⁺ Tex, DCs and CD8⁺ Tn exhibited the largest increase in the M region of the pNR-1 patient (**Fig. 6e**). These findings indicate that immuno-chemotherapy promoted a CD8⁺ T cell–driven anti-tumor TME in the key M region of MPR groups, whereas the pNR-1 patient maintained an immunosuppressive TME, which resulted in tumor cell expansion and T cell exhaustion.

For high-resolution cell–cell and ligand–receptor interactions analysis in intercellular communication, we thus applied 8-μm-spot Decoder-FFPE-seq to explore the cellular neighborhoods of OSCC samples. To ensure methodological robustness, we first benchmarked it against the commercial Xenium platform, confirming its sensitivity and accuracy in cellular-resolution profiling of human clinical samples. We subsetted the post MPR-1 8-μm-spot Decoder-FFPE-seq data from >18,000 genes to the 417 genes shared with the Xenium panel (**Methods and Supplementary Table 14**) to compare gene detection performance in the corresponding common capture regions, respectively (**Supplementary Fig. 16a**).

Comparing the number of UMIs per spot/μm^2^ with the number of transcripts per cell from Xenium, we found that Decoder-FFPE-seq is comparable in sensitivity to Xenium (**Fig. 6f**). Moreover, Decoder-FFPE-seq and Xenium exhibited concordant spatial gene expression and cell-type distribution, exemplified by the tumor-associated epithelial marker *KRT5* and tumor cells (**Fig. 6g and Supplementary Fig. 16b-c**). These results indicate Decoder-FFPE-seq yields remarkable spatial results that comparable to an imaging-based approach.

With 8-μm-spot Decoder-FFPE-seq, we characterized the key M regions of four representative samples from four groups (pre-MPR-1, post-MPR-1, pre-pNR-1, and post-pNR-1, **Supplementary Fig. 17**). Similarly, we identified distinct spatial regions in four samples through unsupervised clustering combined with tumor border annotations from H&E staining, followed by cell-type deconvolution to characterize treatment-induced cellular dynamics (**Supplementary Fig. 18-19**). We obtained cell distributions largely concordant with those derived from 50-μm-spot FFPE-seq, and pre- and post-treatment samples showing consistent cellular dynamic changes (**Fig. 6h and Supplementary Fig. 20a-b**). As in the 50-μm-spot data, CD8⁺ EarlyActive markedly increased in theMPR-1 patient underwent treatment, whereas CD8⁺ Tex expanded in the post-treatment pNR-1 (**Fig. 6h**). We observed that these cells had remarkably spatial interaction pattern changes in the M region before and after treatment (**Fig. 6i**). In MPR-1, CD8⁺ EarlyActive exhibited the highest predictability for the abundance of B cells, CD4 Tn, CD8 Tn, and macrophages before treatment, whereas after treatment they additionally co-localized with B cells, CD8 Tem and Th1. In pNR-1, CD8⁺ Tex shifted from spatial co-localizing with Tfh and CD8 Tem to a broader network including CAFs, CD4 Tn, CD8 Tem, DCs, mast cells, monocytes, plasma cells, Tfh, Th1, and Tregs. These findings suggest that responders effectively transitioned from immune priming to effector maturation during treatment, whereas non-responders developed an immunosuppressive TME that perpetuated T cell exhaustion. Cell–cell communication analysis further confirmed their spatial interactions within the defined M region (**Fig. 6j**). In MPR-1, CD8⁺ EarlyActive interacted robustly with both B cells and Th1 cells via ligand–receptor pairs such as SEMA4D–PLXNB2, SELPLG–SELL, and CCL5–CCR4, suggesting enhanced CD8⁺T cell recruitment and activation. In contrast, pNR-1 was characterized by predominant interactions of CD8⁺ exhausted T cells with CAFs (via FN1–CD44, COL1A1–CD44, COL1A2−SDC4), reflecting a CAF-dominated immunosuppressive microenvironment. Overall. these findings may account for the divergent CD8⁺ T cell dynamics observed between the response groups and highlight their role in shaping distinct therapeutic outcomes.

## Discussion

ST has transformed our ability to map molecular and cellular organization within their original tissue contexts, offering powerful new avenues for both basic biology and clinical pathology. However, most ST methods depend on high RNA integrity (RIN > 7) to obtain high-quality data and are therefore largely restricted to FF material, which requires stringent cold-chain storage and often lacks long-term clinical follow-up. By contrast, FFPE repositories are stable at room temperature for decades, and linked to rich clinical metadata, but their heavily crosslinked and fragmented RNA has limited ST adoption. Existing FFPE-compatible ST approaches, therefore, force trade-offs among throughput, sensitivity, spatial resolution, and accessibility.

Decoder-FFPE-seq addresses these constraints by integrating specific RNA-DNA probe conversion, high-efficient blotting-assisted spatial barcoding, and high-throughput sequencing into a standardized, open workflow. This combination enables genome-wide, spatial cellular profiling of archival tissues while remaining implementable with common laboratory equipment and reagents.

Decoder-FFPE-seq retrievals spatial RNA data with remarkable sensitivity and genome-wide coverage. The split-probe design targets multiple non–poly(A) regions per transcript, converting short RNA fragments into amplifiable DNA probes while minimizing capture of abundant, low-information rRNA (an important limitation of random-priming or in-situ polyadenylation strategies). Paired with blotting onto our nanostructured Decoder slides, which provide ∼10-fold higher barcode density than comparable arrays, this approach prevents barcode loss during harsh FFPE pretreatment and maximizes productive encounters between probes and spatial barcodes. Decoder-FFPE-seq detects ∼617 genes per 8 μm spot (approximately 2.54-4.25 times higher than similar assays), enabling reliable mapping of low-abundance GPCRs and sparse cell populations in the mouse brain.

Beyond polyadenylated mRNAs, Decoder-FFPE-seq extends ST to non-poly(A)-tailed regulatory RNAs and non-host transcripts. Using tailored probe panels, we jointly profiled mRNAs, OSCC-associated lncRNAs, and HPV E6/E7 transcripts in archival tumors. This revealed spatial co-expression modules, such as SM15, in which *NEAT1* acts as a network hub linked to inflammatory mediators and oxidative stress genes. Our approach also showed that local viral RNA abundance correlates with the activation of host proliferation programs. These results demonstrate how RNA profiling in FFPE tissue can reveal multi-layered regulatory that were previously challenging in transcriptome-wide spatial assays.

Decoder-FFPE-seq provides an accessible platform for single-cell–resolved ST studies, lowering technical barriers for a wider community of researchers. Spatially barcoded array-based sequencing methods are particularly attractive because they utilize standard laboratory reagents and instruments, while offering streamlined and time-efficient workflows. Yet, affordable barcode slide fabrication and open-source protocols remain scarce, limiting widespread adoption of FFPE-compatible, single-cell ST. To bridge this gap, Decoder-FFPE-seq combines 1) high resolution Decoder slides with 2) an open-source ST workflow against FFPE samples. Decoder slides, fabricated through a microfluidics-assisted combinatorial barcoding strategy, can be produced cost-effectively in standard laboratory settings following our published protocol. This bypasses the substantial technical hurdles posed by sequencing-based decoding in BGI’s Stereo-seqV2 and the light-directed synthesis required for 10× Genomics’ Visium HD platform. Additionally, we provide a comprehensive, step-by-step FFPE tissue based-ST protocol in the **Supplementary Information**, which covers tissue pretreatment, probe hybridization, and blotting-assisted spatial barcoding. By making both slides fabrication and experimental workflows openly available, Decoder-FFPE-seq ensures broad accessibility across research communities.

Decoder-FFPE-seq unlocks the vast potential of archived FFPE collections for clinical and translational research. FFPE repositories, preserved for decades, hold unparalleled value: they are linked to detailed clinical records, encompass long-term outcomes, and include rare diseases and diverse pathological subtypes. However, RNA in these samples further deteriorates over time, complicating spatial transcriptome recovery and reducing data fidelity. Decoder-FFPE-seq overcomes these barriers. For the first time, we achieved spatial profiling in 20-year-old clinical samples, successfully recovering highly fragmented RNAs (DV200 = 17%) to identify immune populations and detect low-abundance checkpoints. In a 6-year-old OSCC specimen, the method reconstructed TME architecture and key invasion drivers.

Furthermore, in an ICB-treated trial cohort, Decoder-FFPE-seq resolved response-dependent TME heterogeneity and tracked spatial cellular dynamics before and after immunochemotherapy. These findings demonstrate the power of Decoder-FFPE-seq to extract meaningful biological and clinical insights from deeply archived materials.

In summary, Decoder-FFPE-seq establishes a sensitive, genome-wide, single-cell–resolved, and broadly accessible ST framework for spatial biology studies. Looking forward, several refinements could further enhance its capabilities.

Sensitivity may be increased through improved probe release and interfacial barcoding efficiency, such as electrophoretic probe release or DNA nanoengineering approaches to optimize barcode density and orientation^44, 45^. Subcellular resolution may be attainable by integrating high-resolution barcode slides (e.g., nanopatterned arrays)^46–48^ or expansion microscopy into the blotting-assisted workflow^49, 50^.

Expanding probe design to additional RNA species, including microRNAs, other noncoding RNAs, and non-host transcripts, or applying total-transcriptome strategies (e.g., *in situ* polyadenylation, random primer capture), will broaden molecular coverage. Beyond transcriptomics, Decoder-FFPE-seq holds significant potential for spatial multi-omics by incorporating proteomic or epigenomic tagging methods, enabling co-mapping of transcriptome, proteome, and epigenome within the same section^51–53^. Clinically, scaling the approach across large patient cohorts and coupling results with artificial intelligence frameworks will facilitate the discovery of biomarkers, therapeutic targets, and predictive models^54^. Together, these advancements position Decoder-FFPE-seq as a versatile foundation for next-generation spatial omics in biological research and precision medicine.

## Acknowledgments

We thank Jinbang Li, Dunhuang Pan and zhikang Pan for their assistance in bioinformatics analysis. Xiangqian Zhang for the design of the IncRNA probe library. C.Y. was supported by the National Natural Science Foundation of China (22293031, 22521102), Innovative research team of high-level local universities in Shanghai (SHSMU-ZLCX20212601). L.W. was supported by the National Natural Science Foundation of China (22422409, 82341023, and 82227801) and Shanghai Rising-Star Program (23QA1408200).

## Author Contributions

C.Y. conceived the project. C.Y., J.C., and W.L. designed the experiments. J.Z., J.C., and X.F. performed the experiments. Y.M. and Y.G. conducted the bioinformatics analysis. J.C., Y.M., J.Z. Y.L., and Z.Z. analyzed the data. S. H., W.S., and J.Z. prepared the microfluidic chips. J.Z., J.F., and X.X. collected the biopsy samples. J.C. and W.L. wrote the manuscript. C.Y., W.L., and G.C. revised the manuscript and supervised the project.

## Methods

### Reagents

All reagents used in the study are listed in Table 1

**Table 1.**
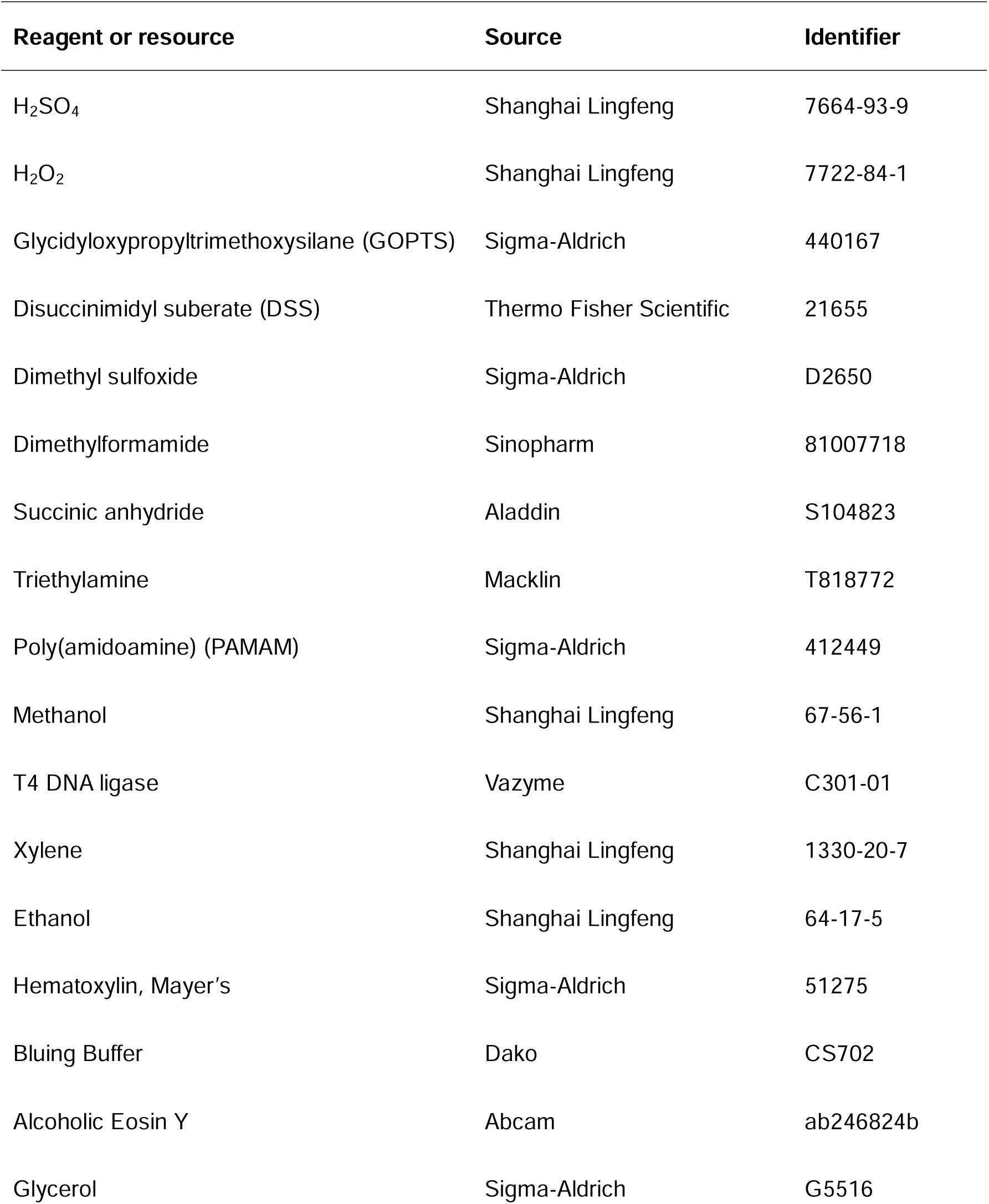
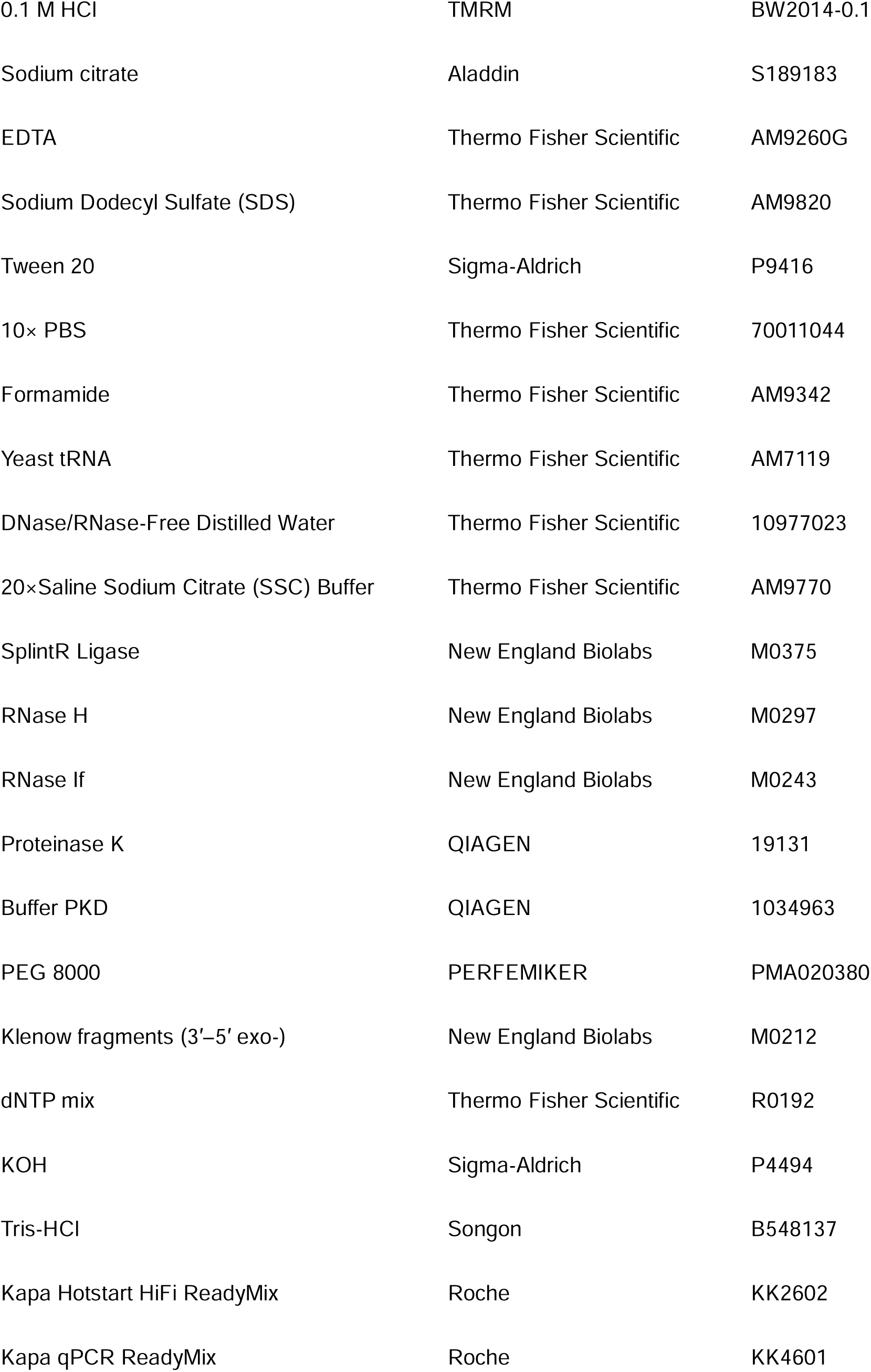
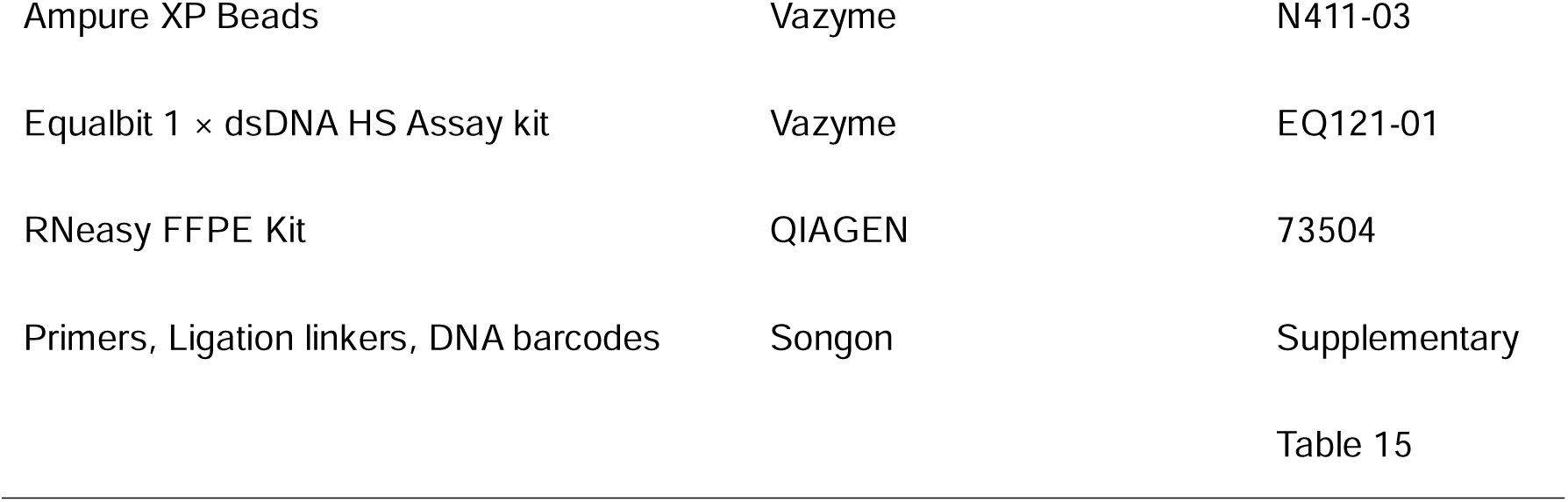
Study reagents.

### Fabrication of Decoder slides

The comprehensive fabrication process, which employs microfluidic-assisted spatial barcoding on 3D dendrimeric slides, has been detailed in our previous publication^19^. Three types of microfluidic chips with different widths and channel numbers were designed: 50 channels*50 μm, 96 channels*15 μm, and 192 channels*8 μm.

Microfluidic chips were fabricated using standard soft lithography. Chrome photomasks (CChip Scientific Instrument Co., Ltd, Suzhou, China) were used to pattern SU-8 photoresist on a silicon wafer, creating the mold. Polydimethylsiloxane (PDMS) pre-polymer (curing agent A: base = 1:10) was poured onto the mold, degassed, cured (120°C, 20 min), peeled, and cut. 1 mm metal punch pen was employed to create inlet/outlet ports. 3D dendrimeric slides were prepared through sequential surface modifications: (1) piranha treatment (2 h) for hydroxyl activation, (2) GOPTS silanization (5 h), (3) blocking with succinic anhydride/triethylamine (1 h), and (4) overnight PAMAM modification. For spatial barcoding, two microchannel chips were sequentially aligned over 3D dendrimeric slides perpendicular with each other. DNA barcode X (5′-aminated) was covalently immobilized via DSS crosslinking, while DNA barcode Y was enzymatically ligated to X using T4 DNA ligase. This generated spatially defined X_i_Y_j_ sequences at channel intersections.

### Design of probe libraries

Probe libraries targeting protein-coding genes of mouse and human transcriptomes were designed with reference to the 10X Genomics (https://www.10xgenomics.com/support/cn/cytassist-spatial-gene-expression/docum entation/steps/probe-sets/visium-ffpe-probe-sets-overview). The mouse probe library has 19,763 probe pairs for 19,465 protein-coding genes (**Supplementary Table 1**). The human probe libraries contain: 1)18,630 probe pairs for 17,943 protein-coding genes; 2) 53,504 probe pairs for 18,085 genes (**Supplementary Table 6**).

We downloaded 333 OSCC-associated lncRNAs from the LncRNA and Disease Database (<u>LncRNADisease3</u>). Using the open-source software Oligo Designer Toolsuite (https://github.com/HelmholtzAI-Consultants-Munich/oligo-designer-toolsuite), corresponding lncRNA probes were designed in three steps: generation, filtering, and probe set selection for each gene. High-quality probes were successfully generated for 275 lncRNA genes (**Supplementary Table 8**). Detailed steps and parameters are provided in the **Supplementary Information**

For probes targeting E6 and E7 mRNAs of high-risk HPV types 16 and 18, we designed them by extending the original 16-nt probes to 25-nt ones based on reported HPV mRNA probes^55^ (**Supplementary Table 8**). Probe filtering was conducted as above-mentioned procedure.

All probes targeting mRNAs, lncRNAs, and HPV 16/18 were synthesized by Beijing Tsingke Biotech Co., Ltd. (Shanghai, China).

### Design of the probe-blotting instrument

The probe-blotting instrument was home-made to achieve precise alignment of tissue sections with Decoder slides, enabling probe transfer and spatial barcoding. It integrates imaging-guided positioning via a camera-equipped, freely moving base for flexible region-of-interest selection, with programmable high-precision temperature and time control, and a chip holder structure. The chip holder structure secures tissue slides on the upper cover while anchoring Decoder slides on the base. A 10-μm-thick film compartment was pasted around the barcoded DNA array on the slide to hold reagents.

### Collection and preparation of FFPE mouse tissues

The whole brain was harvested from euthanizing male C57BL/6J mice (6-8 weeks old; Shanghai JieSiJie Laboratory Animal Co., Ltd.), rinsed with PBS, and dissected into two regions: the MOB and the main brain. All tissues were fixed in 4% paraformaldehyde for 24 h, and embedded in low temperature melting paraffin.

Animal procedures complied with ethical regulations approved by the Laboratory Animal Welfare Ethics Committee (ACU-0501).

### Collection of FFPE patient specimens

Clinically FFPE tissue blocks, originally collected by physicians for diagnostic purposes, were sourced from Department of pathology, Renji Hospital Affiliated to Shanghai Jiao Tong University School of Medicine with the approval of its Ethics Committee (KY2023-013-B), and School and Hospital of Stomatology, Wuhan University, with the approval of the Ethics Committee (WDKQ2025.B42, [2020] Ethics No. 2). Each donor provided written informed consent before participation. Except the breast cancer block, all other FFPE blocks were obtained from School and Hospital of Stomatology, Wuhan University.

OSCC samples encompassed two distinct cohorts: 1) 9 clinically archival OSCC samples stored for different years, including one HPV-positive case, which were sourced from Department of Oral Histopathology, and 2) 12 FFPE specimen blocks derived from seven ICB-treated patients as a subgroup of 68 enrollees in the two-arm, randomized phase II trial (NCT04649476). These samples were sourced from Department of Oral and Maxillofacial Surgery. All the patients in this cohort received neoadjuvant therapy with three cycles of Camrelizumab with or without two cycles of TPF chemotherapy (docetaxel, cisplatin, 5-fluorouracil), followed by curative surgery and adjuvant therapy. From these patients, one non-metastatic lymph node from neck dissection, seven pre-treatment biopsies, and four paired post-treatment samples from surgically resected tissues were obtained.

### Decoder-FFPE-seq protocol

A detailed protocol is provided in the **Supplementary Information**.

### Evaluation of the effect of tissue pretreatment and probe labeling on Decoder slides

To study the impact of key tissue pretreatment and probe labeling on spatial barcodes, Decoder slides were treated as follows: 1) baked at 42°C for 3 h, then 60°C for 2 h; 2) decross-linked at 95°C for 1 h; 3) MOB-mounted Decoder slides had methanol fixation, Tween-20 permeabilization, overnight probe hybridization, and tissue removal. After treatment, Cy3-labelled poly(A) DNA hybridized with treated (n = 3) and untreated slides (n = 3) at 37°C for 30 min, then washed thoroughly. Finally, the slides were imaged and their fluorescence intensity was compared between the treated and untreated control groups (**Supplementary Fig. 2**).

### Immunofluorescence staining and smFISH

Multiplex immunofluorescence staining of the FFPE section from a 6-year-old OSCC sample were performed by Shanghai Ruiyu Biotechnology Co., Ltd., and the slide was imaged and processed using the 3DHistech fluorescence imaging scanner.

smFISH was performed with the SEERNA® RNA FISH Detection Kit (Dynamic Biosystems, China) according to the manufacturer protocol, and the data were processed according to a previously reported study^56^. The target lncRNA (*MALAT1*, *NEAT1*, *H19*) sequences for ISH are shown in **Supplementary Table 9**.

ISH assay for HPV E6/E7 mRNA was conducted on HPV-positive adjacent tissue sections using the HPV detection kit (Dynamic Biosystems, China) according to the manufacturer protocol. For localization analysis, the DAB-stained channels were isolated through RGB-to-HED color space conversion. Gaussian-like signals in the DAB channel images were detected using the Laplacian of Gaussian operator, and the two-dimensional coordinates of all signal points were output. For quantitative analysis, six matched tissue regions were selected, and signal counts within each region were computed individually.

### Xenium in situ high-plex assay

We designed a custom 50-gene panel targeting key markers of CAFs (e.g., collagen genes, FAP, POSTN), tumor cells (e.g., keratins, cadherins, EMT markers), and macrophages (e.g., SPP1, immune-related HLA genes), among others. This panel was integrated into the Xenium Human Immuno-Oncology Profiling Panel, resulting in a final 430-gene panel for the Xenium in situ high-plex assay (**Supplementary Table 14**). FFPE tissue sections (5 μm thickness) were mounted onto Xenium slides and processed according to the manufacturer’s protocol (<u>Xenium In Situ for FFPE – Tissue Preparation Handbook | 10x Genomics</u>). Cellular segmentation was performed based on DAPI-stained nuclear morphology visualized in whole-slide scans acquired using the Xenium analyzer. Analyzer-generated output files were subsequently used for downstream data analyses.

### Data preprocessing of Decoder-FFPE-seq

DynamicST (v1.0.7) was developed to process Decoder-FFPE-seq sequencing data. For read 1, only barcodes x and y with a Levenshtein distance ≤ 1 from the whitelist were retained. UMIs sharing the same (x, y) barcode were clustered using Starcode-umi, and the relationships among read IDs, barcodes, and UMI clusters were recorded. Based on this mapping, a gene expression matrix of UMI counts per gene for each spatial barcode (x, y) was generated for downstream analyses. Transcriptome references included Mus musculus GRCm38 (mm10) and Homo sapiens GRCh38 (hg38), with probe set references “mouse” and “human”.

DynamicST-Assist (v1.2.1) was employed for alignment with H&E-stained histological sections to precisely delineate tissue regions from background areas. Spots located outside the defined tissue boundaries were excluded, along with those exhibiting fewer than 2 UMI counts and 50 detected genes (Note: the gene detection threshold was relaxed from 50 to 25 genes in 8-um spots). Additionally, spots where the percentage of mitochondrial genes exceeded four standard deviations above the mean were removed. All quality control filtering and subsequent SCTransform normalization were performed using Seurat (v5.1.0)^57^.

### Comparison of Decoder-FFPE-seq with similar ST platforms

MOBs, mouse brains, human breast cancer tissue, and lymph nodes were sequenced with Decoder-FFPE-seq, and the results were compared to those obtained from similar ST platforms, including Decoder-seq, 10x Visium Cytassist_FFPE, Patho-DBiT, Visium, Visium HD (8 µm-bin), and Stereo-seqV2 (bin20). Spearman correlation analyses were conducted to assess the degree of similarity between the aggregate expression of each protein-coding gene obtained from Decoder-FFPE-seq and Decoder-seq. The function ‘cor.test’ was used to assess the statistical significance of the correlation coefficient.

### Molecule lateral diffusion analysis

We selected three regional-enriched genes (*Slc17a7*, *Hpca*, and *Ncdn*) to assess RNA lateral diffusion in Decoder-FFPE-seq (8 µm), Slide-seq V2, Visium HD (8 µm-bin), and Stereo-seqV2 (bin20) by reported method^58, 59^. We identified consistent regions (500 µm × 300 µm) exhibiting specific expression of the aforementioned marker genes across these four platforms. Each region was subdivided into six segments (500 µm × 50 µm) for repetition. We summed the UMI counts of each gene at 10 μm intervals (spanning a 50 μm width) along the 500 μm spatial axis. To characterize the distribution of expression, the aggregated expressions were aligned to their spatial coordinates, enabling profiling of peak expression locus and spatial spread. Subsequently, these summed values were normalized and depicted as density curves along the 500 µm axis, with the total area under each curve scaled to unity. Finally, we computed the FWHM of the expression distribution as a metric for transcript diffusion, and compared it to the actual wide of the corresponding region.

### Detection of non-sensory GPCR in mouse brain

The mouse non-sensory GPCR genes were obtained from the ‘Mouse GPCRs + Human and Rat Orthologs’ database^60^ (https://esbl.nhlbi.nih.gov/Databases/GPCRs/), which comprises 790 non-olfactory GPCRs. Genes encoding opsin receptors, taste receptors (TAS1R/TAS2R), vomeronasal receptors, trace amine-associated receptors (TAARs), and Mas-related G protein-coupled receptors were excluded from subsequent analyses. Gene Ontology biological processes enrichment analysis conducted by R package clusterProfiler^61^ was performed to functionally annotate the remaining GPCR genes. The significantly enriched GO terms (adjusted.p < 0.001) were simplified by function ‘clusterProfiler:simplify’ based on the semantic similarity, and were subsequently clustered using function ‘enrichplot:pairwise_termsim’. Lastly, R package ComplexHeatmap was applied for visualization.

To evaluate the performance of Decoder-FFPE-seq in detecting non-sensory GPCR genes, we compared its aggregated expression profile that normalized to average expression per micron with those obtained from Visium HD and Stereo-seqV2. Expression profile consistency between Decoder-FFPE-seq and MERFISH was confirmed by Spearman correlation, with parallel comparisons performed for Visium HD and Stereo-seqV2. Spatial expression patterns of representative GPCR genes were visualized across all four platforms.

### Spatial data clustering and visualization

To detect spatial domains of sequencing sections, we firstly constructed a shared nearest neighbor graph based on the top 50 principal components derived from 2,000 highly variable genes using the ’FindNeighbors’ function. Spatial clustering was then achieved by applying the Louvain algorithm through the ’FindClusters’ function. For visualization, we employed the UMAP algorithm for dimensionality reduction. To identify domain-specific molecular signatures, we conducted Wilcoxon rank-sum tests using the “FindAllMarkers” function to detect differentially expressed genes for each spatial domain. Lastly, domain annotation was accomplished according to their specific expression patterns of the canonical marker and the identified signatures, as well as the pathological annotation on the corresponding H&E sections. Distributions of spatial clusters and gene expression patterns were visualized using Seurat functions, ’SpatialDimPlot’ and ’SpatialFeaturePlot’, respectively.

For the analysis of multiple spatial transcriptomics datasets, particularly pre- and post-treatment OSCC samples, we applied canonical correlation analysis, a built-in Seurat function, to mitigate batch effects and implement data integration within each treatment group (pre- or post-treatment) before above-mentioned clustering analysis.

### Spatial deconvolution of high-resolution mouse brain

We performed cell-type deconvolution on the mouse brain data of 8 µm-spot Decoder-FFPE-seq using Cell2location^62^. The snRNA-seq reference encompassed the mouse whole cortex and hippocampus. The expect average number of cells per spot was set as 2 while other parameters followed the default settings. Following deconvolution, we quantified the cellular composition of each spot, retaining only cell types with predicted abundances exceeding 0.1 for downstream computations.

### Spatial deconvolution and cell-cell colocalization of OSCC

For OSCC datasets, we performed cell-type deconvolution using R package CARD^63^. To construct an adequate scRNA-seq reference dataset, we specifically selected primary tumor specimens from dataset GSE234933^64^. In the dataset, low-quality cells and doublets were filtered out, T cells were annotated by the label ’projecTIL_subtype’ according to the original study, and subtypes of macrophages were classified into macrophages. In each cell population, cells ranking in the top 50% by UMI counts were selected and subsequently down-sampled or up-sampled to 1,000 cells per state, yielding the single-cell expression matrices for 22 cell subtypes.

### Detection of house-keeping genes in data from archival samples

We quantified the expression of six housekeeping genes (*UBC*, *PPIB*, *HPRT1*, *TBP*, *GUSB*, and *ALAS1*) in a 20-year-old OSCC section at pseudo-bulk resolution. Pseudo-bulk samples were generated by applying K-nearest neighbors clustering based on spatial coordinates to group the spots into clusters of 50 spatially adjacent spots each, followed by summation of normalized expression values within each cluster. The pseudo-bulk expression levels of these genes were visualized using box plots and compared against bulk RNA-seq data of primary tumor samples in the TCGA-HNSCC cohort, where the expression were normalized as transcripts per million (TPM).

### Weighted gene co-expression network analysis

We used hdWGCNA^65^, the weighted gene co-expression network analysis tool for sparse data with high dimensions, to investigate the spatial interaction networks between lncRNA and mRNA in OSCC. All expressed features were used to infer the signed networks based on their spatial similarity patterns. Genes related to *NEAT1* was ranked by their connectivity with *NEAT1*, quantified by the topological overlap measure in the common gene module. To validate the investigation of *NEAT1* in OSCC, the standard WGCNA^66^ was applied to TPM-normalized expression matrices of tumor samples in TCGA datasets.

### Colocalization of lncRNA and mRNA

SpaGene^67^ was utilized to evaluate the colocalization of lncRNA and mRNA of interests. Although originally developed to identify spatially variable genes and cell-cell communications, its basic principle is to detect genes colocalized in neighbors. This is achieved by quantifying the spatial connectivity of gene pairs in a subnetwork comprising only spots highly expressing the target genes. Thus, theoretically, SpaGene can estimate the colocalization of any gene pair. Specifically, ‘SpaGene_LR’ function was used to identified if the given lncRNA and mRNA were significantly colocalized, and ‘Plot_LR’ function was used for visualization of the significant colocalization.

### Cell niche identification of OSCC

Spots exhibiting consistent cellular composition patterns across multiple samples were operationally defined as CNs, indicative of conserved structural organization^68^. To systematically identify the CNs representing distinct pathological response groups, we implemented an analytical framework adapted from the literature^68^. First, we calculated the isometric log-ratio (ILR) transformed cellular proportions for each spot. Next, we constructed a SNN graph using ’scran::buildSNNGraph’ function^69^, followed by Leiden clustering to delineate distinct CNs.

### Analysis on tumor margin

To analyze the impact of immuno-chemotherapy on OSCC samples, spatial domains identified through unsupervised clustering (as described in the ’Spatial data clustering and visualization’ section) were pathologically annotated based on corresponding H&E-stained sections. This annotation process categorized tissue regions into four distinct compartments: T, T-M, M-P, and P. We then systematically evaluated: (1) therapy-induced changes within the same compartments, and (2) differences between responsive and non-responsive patients across these four regions, especially the margin sites.

### Test of differential cellular composition

Following the cellular deconvolution, the proportion or abundance of each cell type can be quantified within individual tissue spots. This enabled comparative assessment of cellular composition differences between distinct conditions or regions. Specifically, we identified variations in cell type fraction and determined the overrepresented cell populations within specific tissue niches or regions of interest. This differential analysis was conducted using the Wilcoxon rank-sum test, with statistical significance defined as an adjusted p-value < 0.05 after multiple testing correction.

### Analysis of spatial dependency

CAF-macrophage colocalization of an archival OSCC specimen was quantified using python package DeepCOLOR (https://github.com/kojikoji/deepcolor), which can calculate the colocalization strength of two given cell types in each spot from a single sample. To analyze spatial interactions across multiple samples, R package mistyR^70^

was used to evaluate the spatial dependencies between tumor cells and other cellular components within tumor niches. This approach enabled quantitative characterization of complex spatial dependency among various cell types in TME. Using cell proportion estimates derived from CARD deconvolution across all samples, we constructed multi-view analytical models to quantify interaction patterns. The analysis specifically assessed the significance of spatial interactions between various cell types that located (1) within an individual spot (intra view), (2) between adjacent spots (juxta view), and (3) in a 15-spot radius neighborhood (para view). The median importance scores aggregated across all samples were interpreted as indicators of spatial dependency patterns, revealing either colocalization (positive association) or mutual exclusion (negative association) between cell types under these defined spatial contexts.

### Differential expression analysis

Differentially expressed genes were identified using Seurat’s ‘FindMarkers’ function with the Wilcoxon rank-sum test. Genes meeting the following criteria were considered statistically significant: (1) adjusted p-value < 0.05 and (2) absolute average log2 fold change ≥ 0.5. GOBP enrichment analysis was then performed using R package ’clusterProfiler’ to functionally characterize these differentially expressed genes.

### Gene set enrichment analysis

Single-spot gene set enrichment analysis was perform by R package irGSEA^71^. Using Hallmark gene sets ,GOBP terms and Biocarta pathways from the Molecular Signatures Database^72, 73^ (MsigDB, https://www.gsea-msigdb.org/gsea/msigdb/), tumor-associated gene sets defined by literature^73^. AUCell score was used to quantified the enrichment level. To identify significant differences in gene signature expression between specific niches or regions of interest, “FindMarkers” function was utilized, following the same process described in the “Differential expression analysis” section. To ensure robust and reproducible identification of significantly enriched pathways, ‘irGSEA.integrate’ function was applied to the enrichment scores calculated by AUCell, ssGSEA, UCell and singscores.

### Analysis on colorectal cancer datasets

We analyzed a publicly available spatial transcriptomics dataset (HRA006753) derived from microsatellite instability (MSI) colorectal cancer patients who subsequently responded to immunotherapy^43^. Regions of each section were annotated using aforesaid method (see **Analysis on tumor margin**). Abundances of Immunocytes in margin region were estimated by the average AUCell scores of corresponding cell type marker gene sets.

### Comparison to Xenium

We leveraged SpatialData^74^ to aggregate Xenium transcripts into expression matrices at a single-cell level and to subset the data to align the same regions of 8 µm Decoder-FFPE-seq data. The data from both platforms were preprocessed by retaining only spots/cells that expressed more than 25 genes and genes detected in at least 2 spots/cells, resulting in a final expression profile set of 417 overlapped genes for downstream analysis. We calculated the Spearman correlation coefficient between numbers of UMIs per µm^2^ in Decoder-FFPE-seq and per cell in Xenium within shared genes. Moreover, Cell type prediction was performed separately for spots from Decoder-FFPE-seq and cells from Xenium data using the ‘FindTransferAnchors’ and ‘TransferData’ functions in Seurat. The single-cell expression profile with the common genes from GSE234933, along with its original major cell state annotations, served as the reference. Finally, we visualized the average expression levels of cell type-specific markers across predicted cell types of Decoder-FFPE-seq and Xenium to evaluate concordance.

### Prediction of ligand-receptor interaction

Commot^75^ was utilized to infer ligand–receptor interactions for regions of interests, within a 250 µm distance. Ligand–receptor pairs of cell-cell contact signals, secreted signals and ECM-receptors included in human CellChatDB^76^ were utilized to predict the Ligand–receptor pairs. n analysis on hi-res OSCC sections, each spot was annotated by the top 2 cell types inferred based on deconvolution to predict the potential senders and receivers of significant communication signals. We further investigated whether a positive signal was also enriched in a group of interest (e.g., pNR after treatment). For each significant ligand-receptor pair i in the group k, we defined:

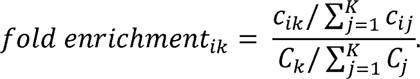

Where *c_ik_* and *C_k_* respectively represent the sum of communication frequency/strength of pair i and all the significant pairs predicted in group *k , K* is the number of groups. We assumed that a ligand-receptor pair is enriched in a specific group if *fold enrichment* > 1.

### Code availability

Toolkits for processing the Decoder-FFPE-seq raw data are open for download from GitHub: https://github.com/DynamicBiosystems/DynamicST and https://github.com/DynamicBiosystems/DynamicST-Assist. Scripts in this study are available in the GitHub repository: https://github.com/miw91/Decoder-FFPE-seq.

### Data availability

All data generated in this study, including raw sequence, expression matrix, space-ranger output and image files will be access under the Gene Expression Omnibus (available upon publication).

